# Macroscopic label-free biomedical imaging with shortwave infrared Raman scattering

**DOI:** 10.1101/2024.06.10.597863

**Authors:** Bernardo A. Arús, Joycelyn Yiu, Jakob G. P. Lingg, Anja Hofmann, Amy R. Fumo, Honglei Ji, Carolin Jethwa, Roy K. Park, James Henderson, Kanuj Mishra, Iuliia Mukha, Andre C. Stiel, Donato Santovito, Christian Weber, Christian Reeps, Maria Rohm, Alexander Bartelt, Tulio A. Valdez, Andriy Chmyrov, Oliver T. Bruns

## Abstract

Shortwave infrared (SWIR) imaging provides enhanced tissue penetration and reduced autofluorescence in clinical and pre-clinical applications. However, existing applications often lack the ability to probe chemical composition and molecular specificity without the need for contrast agents. Here, we present a SWIR imaging approach that visualizes spontaneous Raman scattering with remarkable chemical contrast deep within tissue across large fields of view. Our results demonstrate that Raman scattering overcomes autofluorescence as the predominant source of endogenous tissue background at illumination wavelengths as short as 892 nm. We highlight the versatility of SWIR Raman imaging through *in vivo* monitoring of whole-body tissue composition dynamics and non-invasive detection of fatty liver disease in mice, and identification of calcification and lipids in unfixed human atherosclerotic plaques. Moreover, our approach facilitates the visualization of nerves embedded in fatty tissue, a major advancement for surgical applications. With a simple wide-field setup orthogonal to fluorescence, SWIR Raman imaging holds promise for rapid adoption by clinicians and biologists. This technique opens new possibilities for contrast agent-free visualization of pathophysiology in whole animals and intraoperative imaging in humans.

**Graphical abstract:** 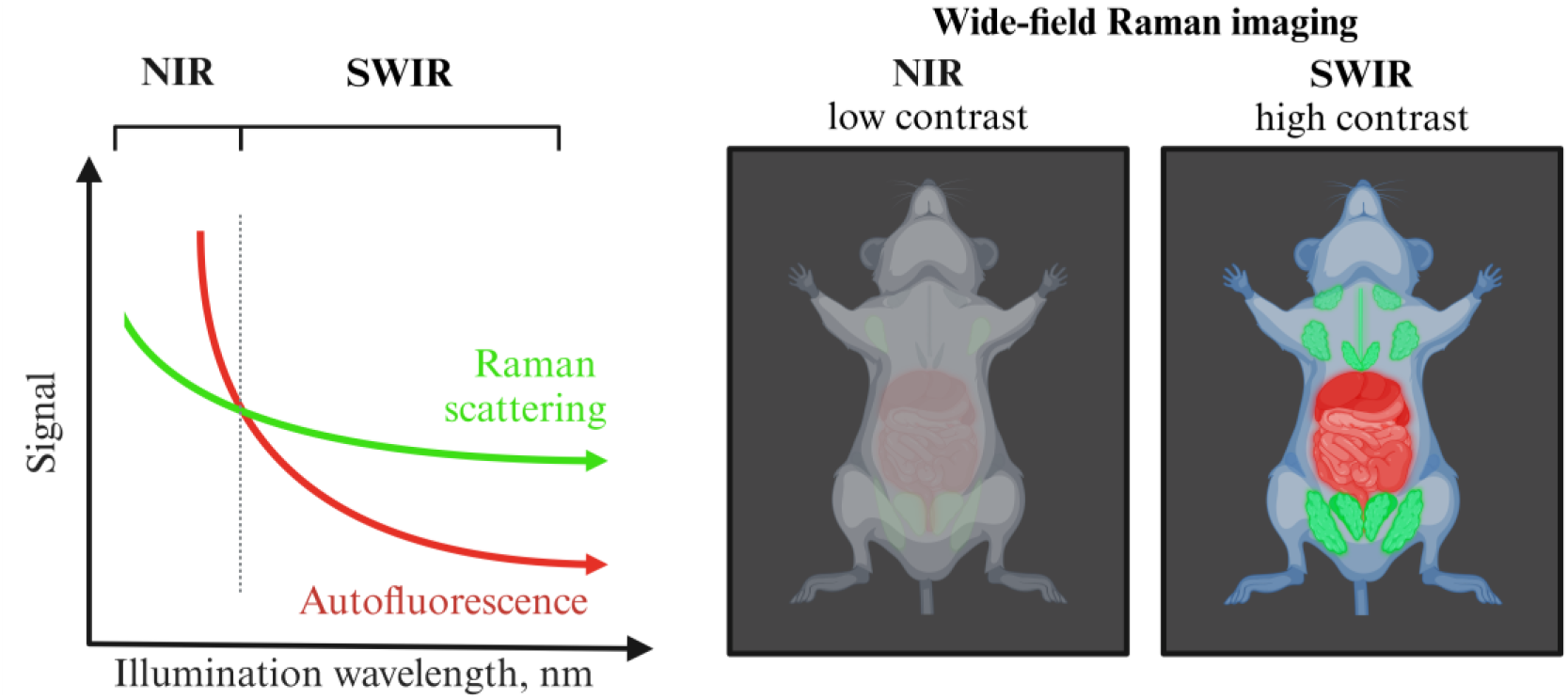

## Introduction

Biomedical imaging in the shortwave infrared (SWIR) range of 1000 to 1700 nm has rapidly expanded over the past decade, primarily due to its significant optical advantages over the visible (400 to 700 nm) and near-infrared (700 to 1000 nm) regions (*1, 2*). This wide adoption was made possible by advances and increased availability of novel SWIR-detecting technology. In biological tissues, SWIR imaging offers deeper penetration, reduced autofluorescence, stronger contrast, and higher resolution compared to other optical windows (*3, 4*). These benefits are particularly evident in animal and clinical studies that compare the SWIR emission tail of indocyanine green, a clinically approved fluorophore, with its near-infrared peak (*5-7*). Contrast agent-based approaches utilize these photophysical properties for a number of applications, including multiplexed fluorescence-guided surgery with tumor delineation (*8*), small metastasis detection (*9*), blood flow mapping (*10, 11*), high-resolution vascular imaging through mouse skin and skull (*12*), and multiplexed tissue imaging in freely moving animals (*13*). Furthermore, SWIR imaging has demonstrated label-free applications by detecting middle ear pathologies (*14, 15*) and cerebrospinal fluid leaks (*16*) based on water absorptivity, and monitoring liver disease through endogenous lipofuscin autofluorescence (*17*).

Raman scattering imaging is a well-established label-free technique used to analyze chemical composition in various materials, including biological tissues (*18, 19*). Particularly by enhancing Raman scattering intensity, these methods have demonstrated remarkable capabilities in molecularly characterizing tissues and cells in both clinical and pre-clinical settings (*20-23*). However, their application has been confined to the microscopic scale (*24*). Expanding Raman imaging to larger fields of view would enable comprehensive analysis of heterogeneous tissues, paving the way for novel applications in whole-body animal studies and medical procedures (*25, 26*). Wide-field Raman imaging has this potential, but conventional near-infrared approaches are hindered by tissue autofluorescence (*24, 27-29*).

Here, we introduce a method that utilizes SWIR imaging to enable spontaneous Raman scattering visualization deep within tissues across large fields. By using illumination wavelengths between 892 and 1064 nm, our approach overcomes previous autofluorescence limitations in spontaneous Raman imaging of biological tissues and offers unprecedented chemical contrast. To demonstrate the power of SWIR Raman imaging, we showcase its ability to non-invasively monitor dynamic physiological and pathological tissue remodeling in mouse models and human tissue.

### Design and *in vitro* validation of wide-field macroscopic SWIR Raman imaging

To image macroscopic spontaneous Raman scattering in the SWIR spectral region, we designed a straightforward imaging system that comprises components similar to those detecting SWIR fluorescence (*30*). Raman imaging was achieved by diffusely illuminating the sample with specific wavelengths and filtering the scattered light with narrow band-pass filters (**Fig. 1 a** and **Table S1-2 a**). These filters selectively captured the scattered wavelengths corresponding to the Raman shifts of interest, which were determined as a combination of the illumination wavelength and the wavelength corresponding to the energy shift of specific bonds, according to

**Figure 1.**
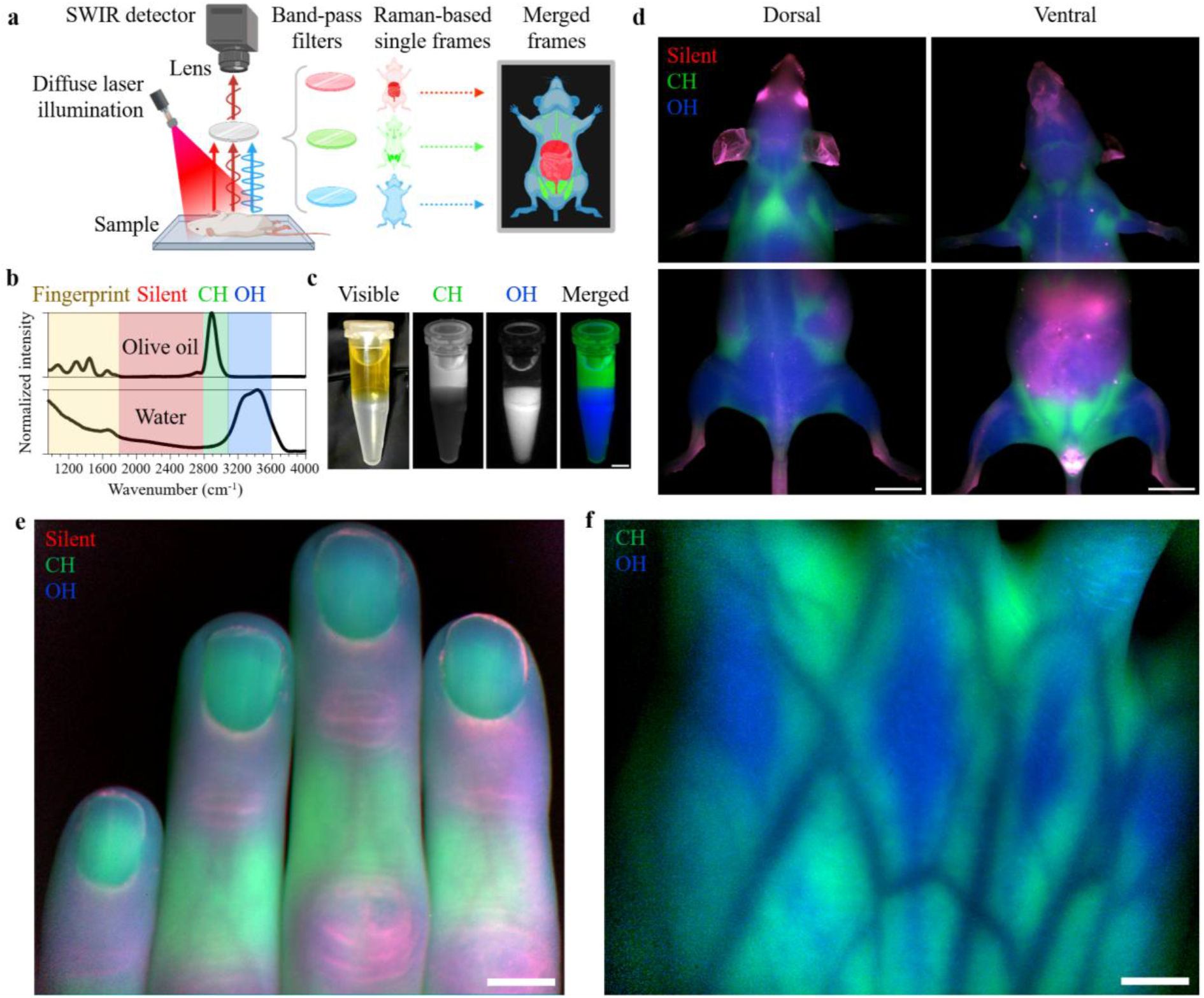
Wide-field macroscopic SWIR Raman imaging system design, *in vitro* validation, and initial biological demonstrations. **a)** Schematic representation of the SWIR Raman imaging setup. Samples are diffusely illuminated with a laser, and the resulting Raman scattering photons pass through specific band-pass filters and arrives at a deep-cooled InGaAs detector sensitive to shortwave infrared (SWIR) wavelengths. Individual bands are acquired sequentially, have pseudocolors assigned, and are displayed merged. **b)** Raman scattering spectra of pure olive oil and water. The four major Raman scattering regions are labeled. **c-f)** Wide-field SWIR Raman imaging of (c) a two-phase mixture of olive oil and water in a 1.5-mL polypropylene tube, (d) an intact healthy C57BL/6J mouse, (e) the dorsal surface of human fingers, and (f) the dorsal surface of the stretched right hand. The Raman-silent, CH and OH regions are shown in red, green or blue, respectively. The displayed contrast was optimized separately for each scattering band; individual scattering bands with corresponding intensity bars are displayed in Fig. S4 and S6. The laser and filter combinations used for each scattering band are listed in Tables S1-2. Scale bars: (c) 0.5 cm; (d)-(f) 1 cm.

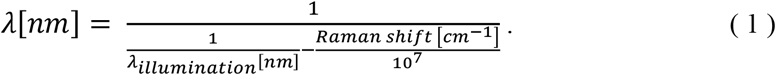

As an example, upon 892 nm illumination, the Raman scattering of the CH region between 2800 and 3100 cm^-1^ shifts to the region between 1189 and 1233 nm, while that of the OH region between 3200 and 3600 cm^-1^ shifts to the region between 1248 and 1314 nm (**Fig. 1b** and **Fig. S1**).

To illustrate and validate the ability of our system to detect specific Raman scattering from different compounds, we imaged water and olive oil, which were spectrally distinguished according to their Raman scattering shifts in the CH and OH regions, respectively (**Fig. 1 b** and **Fig. S1**). By using 892-nm light and sequentially imaging one frame with a narrow band-pass filter selective for the CH region and a second frame with a filter selective for the OH region, we clearly distinguished olive oil and water based on their chemical contrast on a two-phase mixture in a polypropylene tube **(Fig. 1 c**). This approach was extended to other illumination wavelengths and Raman bands, including the fingerprint region, enabling the distinction among hydroxyapatite, proteins and lipids after 1064 nm illumination, based on respective Raman scattering in the PO_4_^-3^ (imaged between 888 and 1065 cm^-1^), amide I (imaged between 1555 and 1706 cm^-1^), and CH region (**Fig. S2-3)**. These findings corroborate that SWIR Raman imaging effectively captures distinct chemical components in a wide-field configuration, akin to a straightforward wide-field fluorescence setup. By selecting specific illumination wavelengths and detection filter combinations, chemical contrast is achieved within single frames, with the only necessary image processing being dark current correction.

### Chemical contrast imaging through skin in whole mice and human hands

Building up on the newly enabled ability to visualize Raman scattering across large fields of view, we investigated the whole-body chemical contrast of biological tissues within intact mice. Adipose tissue fat depots and the xiphoid cartilage, located below the sternum, exhibited strong contrast through intact skin in the CH region, while signal in the OH band was distributed throughout the entire body **(Fig. 1 d and Fig. S4)**. Furthermore, subcutaneous lymph nodes and adipose tissue fat depots were clearly discernible, based on their relative water and lipid content, determined by their respective OH and CH Raman intensities **(Fig. S5)**. This chemical contrast distinctly stood out from background autofluorescence, which we imaged by selecting a filter in the Raman-silent region **(Fig. 1 d and Fig. S4)**. Autofluorescence showed higher signal in skeletal bone, liver, intestine, and skin areas rich in melanin, consistent with previous reports (*31*).

Moreover, SWIR Raman imaging enabled the visualization of chemical contrast through human skin. We visualized subcutaneous fat pads on the dorsal surface of the hand and fingers by their CH Raman signal, while the OH Raman band highlighted the extensor tendons and adjacent dorsal interossei muscles of the hand **(Fig. 1 e-f and Fig. S6)**. Notably, the Raman contrast of fat pads and tendons was observed underneath subcutaneous blood vessels, which yielded negative contrast, particularly in the CH region. In contrast, autofluorescence predominantly manifested in the skin surface creases.

These initial findings highlight the capability of SWIR Raman imaging to non-invasively identify biological tissues based on chemical contrast through skin, across fields of view larger than 50 cm^2^, without contrast agent administration. Notably, these results were obtained with diffuse illumination within the international laser safety limits (*32*), underscoring the compatibility of SWIR Raman imaging with *in vivo* applications in both animal models and humans.

### Dominance of Raman scattering as tissue background at ≥892 nm illumination

To understand the influence of different illumination wavelengths on chemical contrast in tissues and determine their advantages and limitations, we acquired SWIR Raman images from intact mice illuminated from 785 nm, a commonly used wavelength in Raman imaging, to 1064 nm, a red-shifted wavelength that extends Raman scattering wavelengths beyond the limits of standard InGaAs-based detectors. Our findings revealed that the chemical contrast, particularly that of the CH region in adipose tissue fat depots, became stronger than autofluorescence once the illumination wavelength reached 892 nm, and continued to rise up to 1064 nm (**Fig. 2 a-d** and **Fig. S7-8)**. This increase in Raman-based contrast was evident even when using long-pass filters capturing the Raman-silent region within the same frame, highlighting that the enhancement in chemical contrast at longer illumination effectively overcomes the contribution of autofluorescence even within non-specific broad imaging bands (**Fig. S9)**. However, longer wavelengths required slower imaging frame rates (**Table S1)**, emphasizing the need to balance contrast and acquisition speed according to specific application requirements.

**Figure 2.**
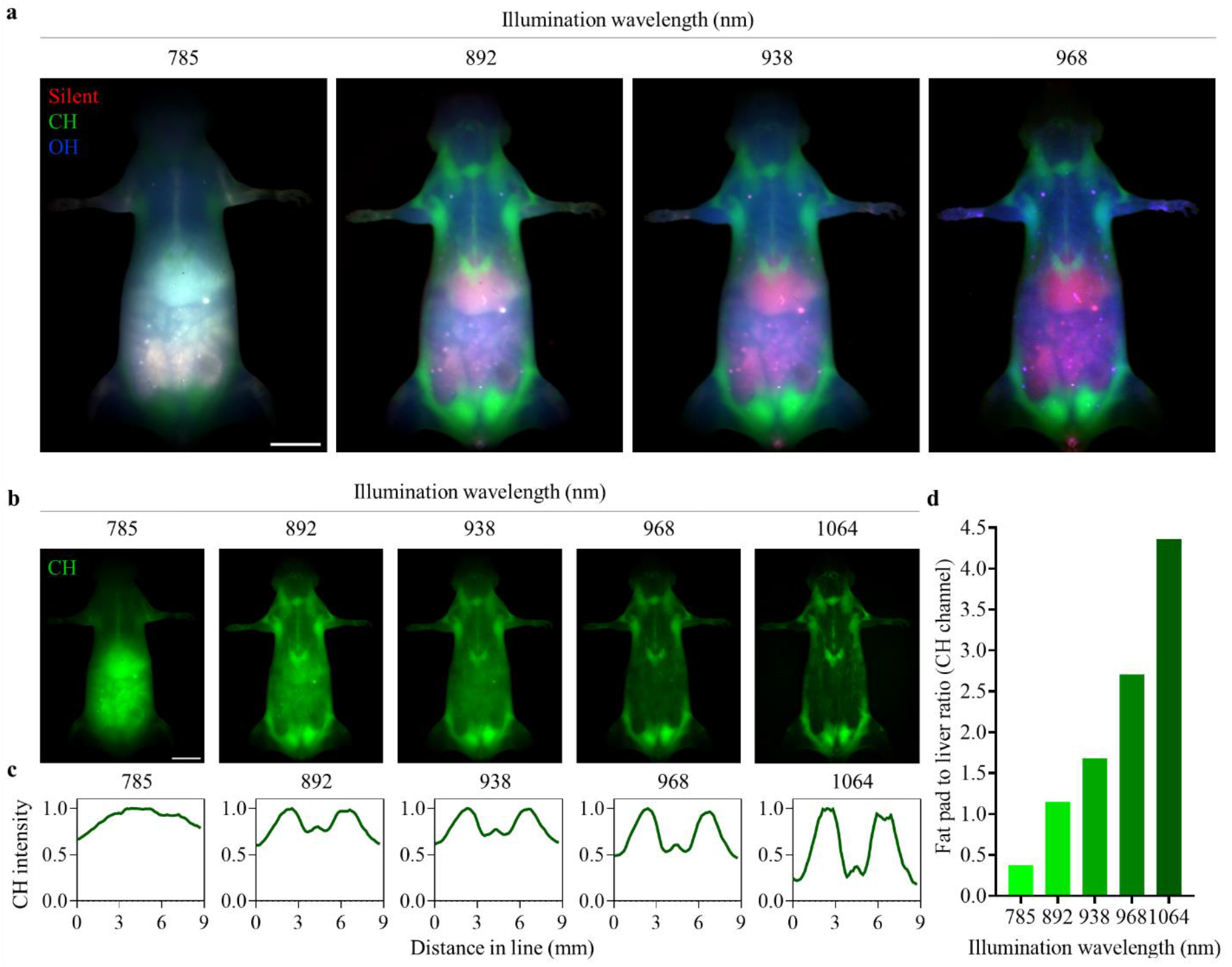
Dominance of Raman scattering as the main source of tissue background in wide-field imaging with illumination wavelengths equal or above 892 nm. **a)** Wide-field SWIR Raman imaging of an intact healthy C57BL/6J mouse at illumination wavelengths of 785, 892, 938 or 968 nm. The Raman-silent, CH and OH regions are displayed in red, green or blue, respectively. **b)** Individual CH region frames for the merged images in panel (a), with the addition of 1064 nm illumination. **c)** Pixel intensity profile along a line crossing the xiphoid cartilage on the mouse chest, extracted from the CH region images in panel (b). **d)** Ratio of mean CH intensity between fat pad and liver, measured for each illumination wavelength, serving as a chemical contrast metric. The displayed contrast was optimized separately for each scattering band. Individual images with corresponding intensity bars are displayed in Fig. S7, while the regions of interest (ROIs) drawn to extract the line profile in panel (c) and the fat pad to liver ratio in panel (d) are displayed in Fig. S8. The laser and filter combinations used for each scattering band are listed in Tables S1-2. Data are representative of two mice. Scale bars: 1 cm.

### Non-invasive SWIR Raman imaging for chemical contrast in mouse disease models

Subsequently, we explored the application of SWIR Raman imaging to non-invasively visualize changes in body composition at the whole-body level in preclinical mouse models. Initially, to detect adipose tissue phenotypes, we imaged genetically obese mice (*ob*/*ob*), which are hyperphagic due to the lack of the satiety hormone leptin, resulting in up to a threefold increase in body weight compared to age-matched controls (*33*). The abundance of adipose tissue fat depots throughout their bodies was highlighted by Raman CH region contrast (**Fig. 3 a** and **Fig. S10)**.

**Figure 3.**
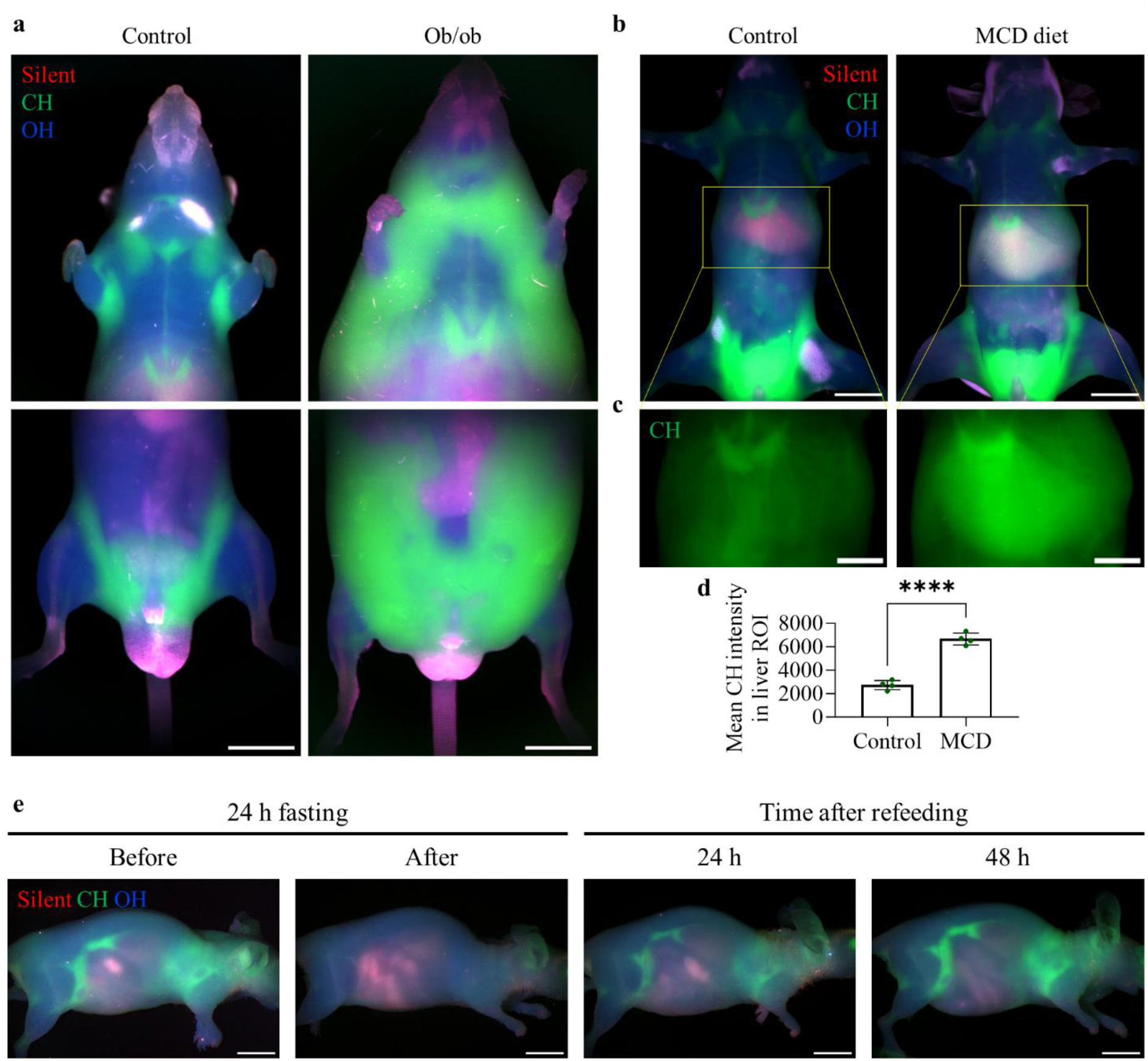
Non-invasive *in vivo* SWIR Raman imaging of chemical contrast in mouse pathophysiology. **a)** SWIR Raman imaging of a genetically obese mouse (ob/ob) and an age-matched lean C57BL/6J control mouse, representative of 2 mice per genotype. **b-c)** SWIR Raman imaging of a C57BL/6N mouse after 6 weeks on a diet deficient in methionine and choline (MCD) or a normal diet (control), representative of 4 mice per group. (b) Whole-body view. (c) Cropped view of the liver region displaying only the CH region. **d)** Quantification of mean CH Raman intensity values (counts/min) in a liver region of interest (ROI) extracted from the MCD and control mice. Differences between the groups were statistically analyzed using an unpaired, two-tailed t test. **** indicates P-value < 0.0001. **e)** *In vivo* imaging of body composition changes in anesthetized nude mice after fasting and refeeding. Animals were imaged at baseline (before), after 24 h of fasting, then after 24 and 48 h of refeeding. Results are representative of two animals. The Raman-silent, CH and OH regions are displayed in red, green or blue, respectively. The displayed contrast was optimized for each scattering band, and, except for panel (a), the same intensity values are displayed for each image within a panel to enable direct comparison. Individual images with corresponding intensity bars are displayed in Fig. S10-S11 and S14, while Fig. S12 displays the biological replicates of panels (b)-(d), and Fig. S13 shows the ROIs drawn to extract the mean values in panel (d). The laser and filter combinations used for each scattering band are listed in S1-2. Scale bars: (a)-(b) 1 cm; (c) 0.5 cm; (e) 1 cm.

Next, we assessed the potential of SWIR Raman imaging for the non-invasive chemical contrast detection of liver steatosis, which is characterized by lipid accumulation in the liver. The label- and contact-free detection of liver steatosis is essential for monitoring liver disease progression, such as metabolic dysfunction-associated steatotic liver disease (MASLD, also called non-alcoholic fatty liver disease, NAFLD), and evaluating treatment effectiveness (*34*). We applied our method to visualize MASLD-associated tissue changes induced by a methionine and choline-deficient (MCD) diet (*35-37*) by conducting macroscopic SWIR Raman imaging of mice following a six-week feeding period. Through intact skin, we detected stronger Raman CH intensity in the livers of the mice fed the MCD diet than in littermates fed a normal diet (**Fig. 3 b-d** and **Fig. S11-13**). These findings demonstrate the potential of SWIR Raman imaging as a valuable tool for whole-body, non-invasive monitoring of tissue composition changes during the progression of diseases such as MASLD.

### *In vivo* monitoring of tissue dynamics with whole-body SWIR Raman imaging in mice

Turning to *in vivo* applications, we utilized SWIR Raman imaging to longitudinally monitor tissue dynamics in living mice, showcasing its non-invasive and label-free properties. We explored the ability of our system to track *in vivo* body composition changes in mice exposed to fasting, a process known to swiftly mobilize adipose tissue fat depots, which typically replenish within 48 hours of refeeding (*38, 39*). Notably, we observed a strong reduction in Raman CH region scattering from adipose tissues following a 24-hour fasting period, which was reversed within 48 h of refeeding (**Fig. 3 e** and **Fig. S14)**.

To expand its utilization for longitudinal visualization of whole-body tissue dynamics, we employed SWIR Raman imaging to monitor tissue modifications during tumor growth. To accomplish this, we tracked the development of 4T1 breast cancer cell xenografts implanted subcutaneously in mice. Imaging conducted over a 9-day period after tumor cell implantation revealed tumor-associated edema, distinguishable by OH contrast against the surrounding CH contrast of adipose tissue (**Fig. S15-16**).

Furthermore, to validate the orthogonality of SWIR Raman imaging in conjunction with traditional fluorescence for *in vivo* studies, we implanted mice with 4T1 cells engineered to express the fluorescent protein iRFP720. While its emission peak is in the near infrared, the emission tail of iRFP720 can be detected in the SWIR (*40*). This dual approach allowed us to monitor both tumor cell fluorescence and whole-body Raman scattering over time (**Fig. S17**). These results demonstrate the versatility of wide-field SWIR Raman imaging for monitoring physiological and pathological changes longitudinally, as well as the potential to combine this technique with fluorescence markers for comprehensive *in vivo* studies.

### SWIR Raman imaging for *ex vivo* analysis of unfixed tissue composition in health and disease

We next aimed to characterize the application of our system to analyze tissue composition in dissected samples *ex vivo*, a process often entailing time-consuming steps like fixation, slicing and staining, or other procedures that may also alter tissue composition (*41, 42*).

Firstly, we dissected organs from healthy mice, and expanded our *in vivo* findings by detecting stronger OH Raman scattering contrast in skeletal muscle, kidneys, and heart, indicating abundant water content, while contrast of the brain was particularly strong in both the CH and OH regions, reflecting the high content of both lipids and water (**Fig. S18**). As expected, different adipose tissue fat depots exhibited high CH contrast, and the inguinal lymph nodes also exhibited chemical contrast with the adipose tissue fat depots in which they were embedded. The autofluorescence signal was consistent with the *in vivo* findings, showing stronger contribution in the liver, intestine, and melanin-rich skin patches. Moreover, SWIR Raman imaging distinguished gray and white matter in the porcine brain, based on their respective OH and CH intensities (**Fig. S19-20**). Interestingly, the sulcus in the cortical regions could be differentiated from the gyrus based on their Amide I contrast (**Fig. S20**).

Turning to pathological conditions, we investigated the application of SWIR Raman for measuring lipid content in unfixed tissues, which would enable direct tissue analysis while preserving its native and intact composition. Indeed, we observed stronger CH intensity, indicative of higher lipid content, in livers from genetically obese mice (**Fig. S21**), as well as from wild-type mice fed either a MCD diet (**Fig. 4 a-c** and **Fig. S22**), or a high-fat diet known to induce obesity and liver steatosis (**Fig. S23 a-b**). Notably, we successfully imaged up to 24 liver samples from different mice in a single frame, demonstrating the high-throughput capability of SWIR Raman for macroscopic detection of lipid content in unprocessed tissues.

**Figure 4.**
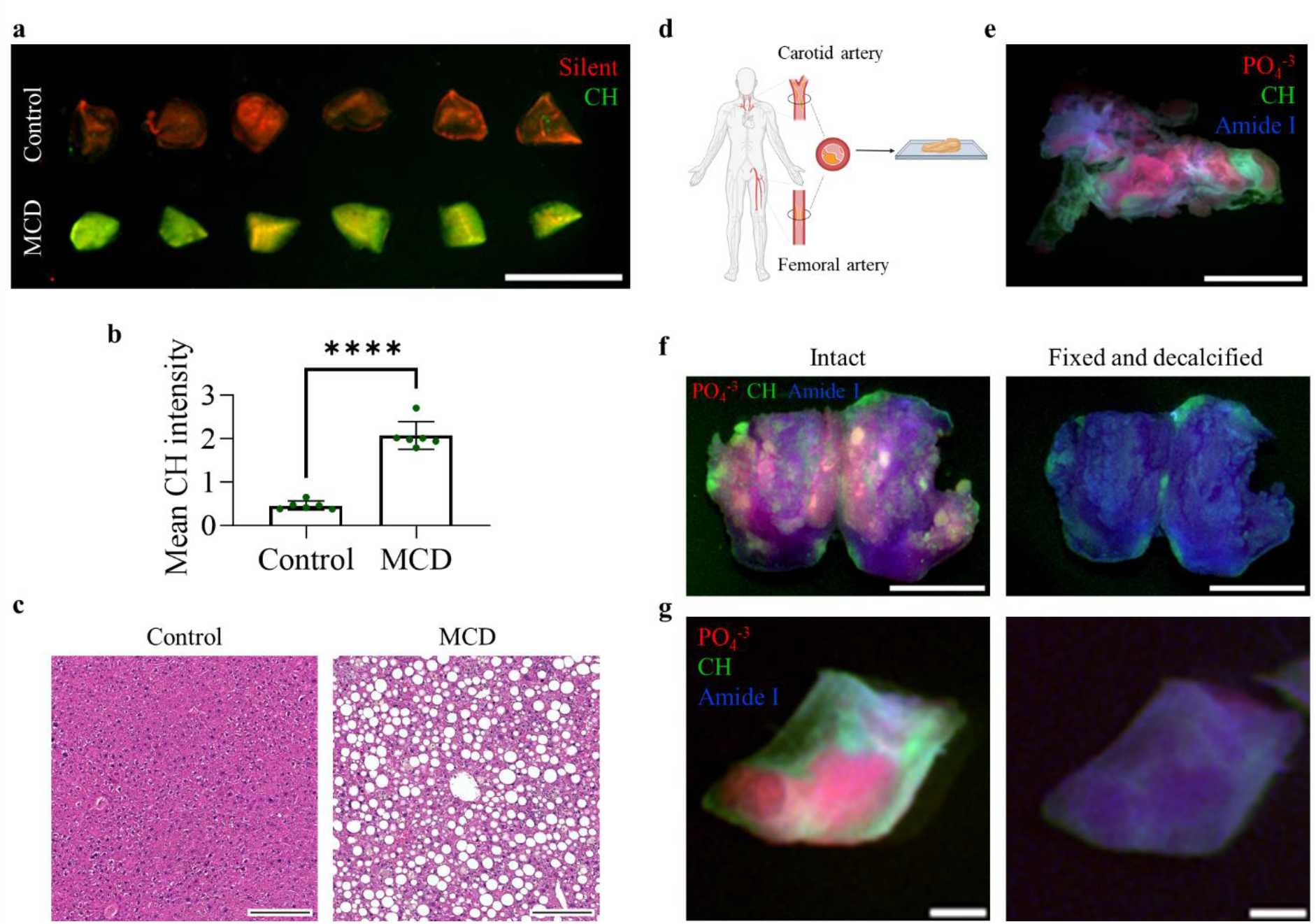
SWIR Raman imaging for *ex vivo* analysis of tissue composition in unfixed samples from mice and humans. **a-c)** Analysis of livers excised from C57BL/6N mice after 7 weeks on a diet deficient in methionine and choline (MCD) or on a normal diet (control). Liver samples were analyzed in terms of (a) SWIR Raman imaging with illumination at 1064 nm, and (b) quantitation of mean CH Raman intensity [counts/min/(mW/cm^2^)] based on the image in panel (a). Differences between the groups were statistically analyzed using an unpaired, two-tailed t test. **** indicates P-value < 0.0001. The Raman-silent and CH regions are displayed in red and green, respectively. (c) Representative histological analysis of the liver samples in panel (a) with hematoxylin and eosin staining. **d-g)** SWIR Raman tissue analysis of human atherosclerotic plaques from carotid and femoral artery surgeries. (d) Schematic representation of the surgical biopsy. The plaque is removed from the artery, and the intact specimen is subjected to imaging. (e) Atherosclerotic plaque excised from a human femoral artery, imaged in the SWIR Raman imaging regions corresponding to calcification-associated PO_4_^-3^ (red), protein-associated amide I (blue) and lipid-associated CH (green). (f)-(g) Comparison between Raman images of intact atherosclerotic plaque specimens (left) and after a fixation and decalcification protocol (right) of a clinical specimen from a (f) femoral artery and (g) a carotid artery. The displayed contrast was optimized for each scattering band; in panels (f)-(g), both frames within each panel display the same intensity values to enable direct comparison. Individual images with corresponding intensity bars are displayed in Fig. S22 and S24-S25. The laser and filter combinations used for each scattering band are listed in Tables S1-2. Scale bars: (a) 1 cm; (c) 200 µm; (e)-(f) 1 cm; (g) 0.2 cm.

Atherosclerosis is an inflammatory disease prompted by abnormalities in lipoprotein metabolism and characterized by accumulation of lipids and inflammatory cells in the vessel wall (*43-45*). Analyzing atherosclerotic plaques presents challenges, as important components for atherosclerosis scoring and risk stratification, such as calcified areas and accumulated lipids, can be lost during sample preparation for traditional histological analysis (*46, 47*). To complement microscopic histological assessment, we employed SWIR Raman imaging to visualize heterogeneous tissue composition in unfixed human plaques. Utilizing excised atherosclerotic tissue from carotid or femoral arteries obtained from patients during eversion thromboendarterectomy, we targeted the PO_4_^-3^ Raman peak at 960 cm^-1^ to detect atherosclerotic plaque calcification and the CH region to detect lipids (*48, 49*). Notably, we observed distinct domains showing calcification or substantial lipid content within plaques, without the need for tissue processing, such as fixation, decalcification, slicing and staining (**Fig. 4 d-e** and **Fig. S24**). We validated the specificity of Raman contrast for calcification by comparing imaging results before and after employing a fixation and decalcification protocol. This protocol eliminated the PO_4_^-3^ signal and reduced the CH intensity (**Fig. 4 f-g** and **Fig. S25**). Moreover, the comparison between imaging of the whole biopsy vs. common histological sections revealed the insufficiency of traditional methods to capture the heterogeneity of atherosclerotic plaque composition (**Fig. S26)**.

In conclusion, our results highlight the effectiveness of SWIR Raman imaging for analyzing tissue composition in large and heterogeneous samples *ex vivo*, without the need for tissue processing while preserving its native composition.

### Nerve identification in a surgical model using SWIR Raman imaging

The precise identification and preservation of nerves pose major challenges in surgical procedures, as nerve misidentification or damage can lead to severe complications such as paralysis, incontinence, or erectile dysfunction (*50, 51*). While nerves can be visually differentiated from muscle tissue, their identification under visible light becomes very challenging when they are embedded in fatty tissue (**Fig. 5 a)**. We employed SWIR Raman imaging to visualize the facial nerve in a porcine model. Our results showed strong nerve contrast within fatty tissue based on Raman imaging (**Fig. 5 b-c and Fig. S27)**. Most importantly, SWIR Raman imaging was able to reveal hidden structures otherwise invisible under visible light. Despite the presence of the OH signal in blood vessels, which can be visually identified, our findings demonstrate the potential of SWIR Raman imaging to be employed in intraoperative settings. By utilizing label-free chemical contrast, it enables the visualization of critical structures like nerves without the need for exogenous contrast agents.

**Figure 5.**
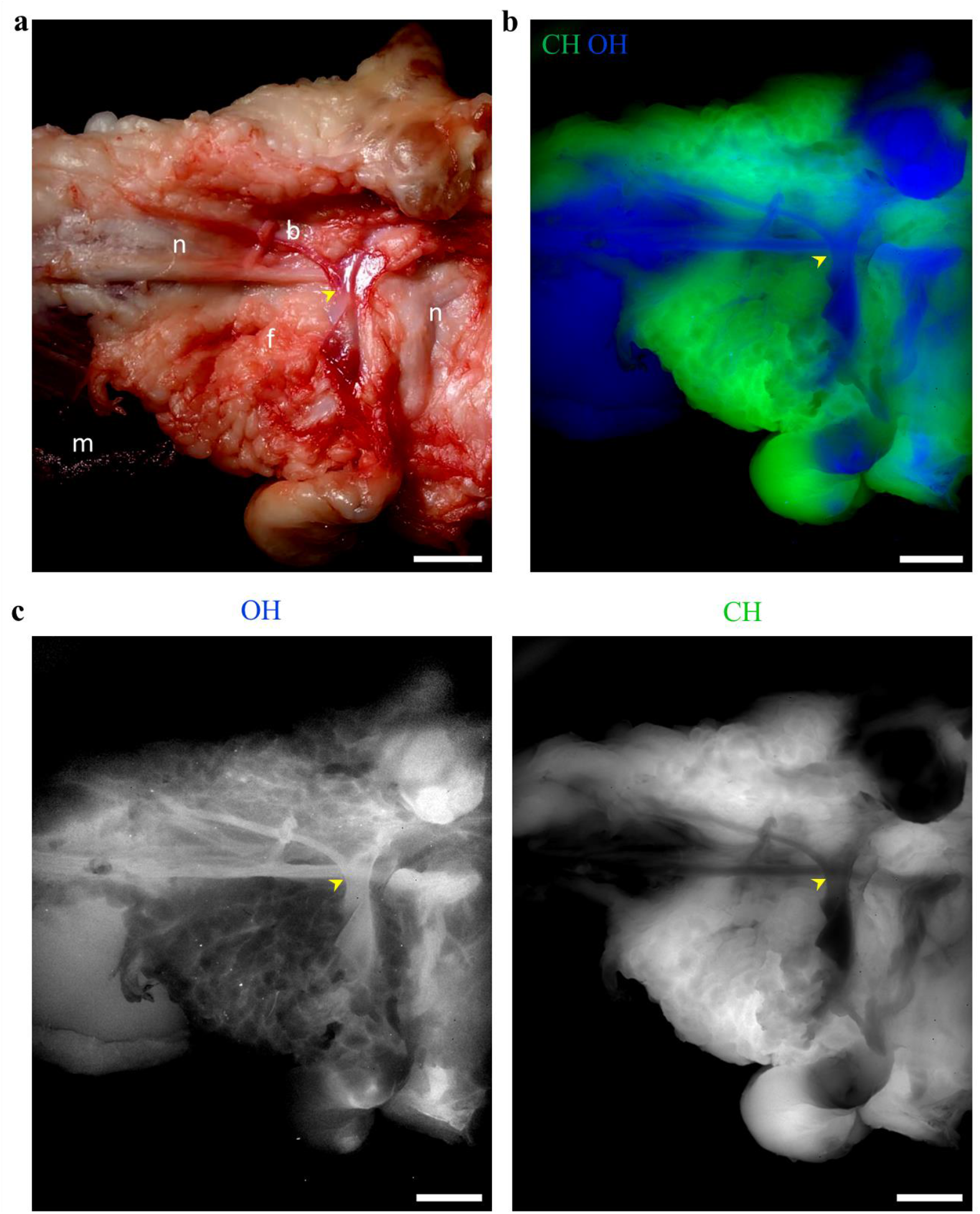
Nerve identification in fatty tissue of a porcine surgical model using SWIR Raman imaging. **a-c)** A porcine facial nerve was exposed, and imaged (a) in the visible and (b-c) using SWIR Raman imaging. The contrast of facial nerve to fatty tissue was obtained using the OH (blue) and CH (green) Raman imaging regions, revealing hidden parts of the nerve (yellow arrow head). The displayed contrast was optimized separately for each scattering band; individual scattering bands with corresponding intensity bars are displayed in Fig. S27. The laser and filter combinations used for each scattering band are listed in Tables S1-2. Legend: facial nerve (n), blood vessel (b), muscle (m), fatty tissue (f). Scale bars: 1 cm.

## Discussion

In this work, we unveil a novel Raman scattering-based imaging method that enables novel biomedical applications through its ability to achieve high chemical contrast in single frames without autofluorescence subtraction or spectral unmixing, with fields of view surpassing 50 cm^2^. Our findings reveal that Raman scattering-based chemical contrast overcomes autofluorescence when tissues are illuminated with wavelengths of 892 and longer. This suggests that wide-field Raman imaging should employ these wavelengths for optimal chemical contrast, thereby requiring SWIR Raman scattering detection. In addition to autofluorescence suppression, SWIR Raman imaging capitalizes on the optical benefits inherent to this wavelength range, such as deeper tissue penetration with stronger imaging contrast and higher resolution (*2, 3*). While it is conventionally conceived that spontaneous Raman detection requires long exposure times (*24, 52*), our results show distinct chemical contrast with high signal to noise within a 10-second exposure per frame under illumination at 892 nm. Employing larger and more sensitive SWIR detectors, which are currently being developed, will enable a higher signal-to-noise ratio and shorter exposure times.

The ability of SWIR Raman to image live and unfixed tissues with chemical contrast has significant implications to both pre-clinical and clinical settings. Compared to conventional modalities, like magnetic resonance imaging (MRI), computer tomography (CT), and ultrasound (*53-55*), SWIR Raman imaging offers distinctive advantages. CT and ultrasound provide anatomical information but lack chemical contrast. In turn, MRI can provide similar contrast but requires complex infrastructure. SWIR Raman setups can be compact and mobile, utilizing a simple optical configuration akin to fluorescence imaging. These characteristics will facilitate the implementation of SWIR imaging in peri- and intraoperative settings in the future.

In conclusion, SWIR Raman imaging not only broadens the applications of Raman imaging to the macroscopic scale, but also expands the overall capabilities of biomedical imaging. By remaining orthogonal to classical fluorescence imaging, this straightforward approach supports multimodal strategies, thereby opening new avenues for pre-clinical and clinical applications. In surgical settings, its use promises immediate benefits by enabling the visualization of critical structures, such as nerves concealed within fatty tissue, thereby addressing a major risk faced by patients undergoing surgery today.

## Acknowledgments

The authors thank Thomas Bischof for critical discussions related to this manuscript, Martin Warmer for help in setting up hardware, Hannes Rolbieski for assistance with animal handling and tumor cell culture, Julia Barthel for contributing with the facial nerve dissection, and Pamela Sabarstinski for processing of arterial biopsies. The authors also thank the Core Facility Pathology & Tissue Analytics at Helmholtz Munich for performing the histological analysis.

## Funding

Helmholtz Zentrum München core funding (O.T.B)

National Center for Tumor Diseases (NCT) core funding (O.T.B)

Deutsche Forschungsgemeinschaft (DFG) Emmy Noether program no. BR 5355/2-1 (O.T.B)

German Federal Ministry of Education and Research (BMBF) project BetterView (O.T.B)

National Center for Tumor Diseases (NCT) PoC grant project SATISFIES (O.T.B)

Helmholtz Imaging Project grant ZT-I-PF-4-038 (O.T.B)

Chan Zuckerberg Initiative (CZI) Deep Tissue Imaging grants DTI0000000248 and DTI2-0000000206 (O.T.B, A.C.)

Deutsche Forschungsgemeinschaft (DFG) SFB1123-B10 (O.T.B., A.B.)

Deutsche Forschungsgemeinschaft (DFG) SPP2306 BA4925/2-1 (A.B.)

Deutsches Zentrum für Herz-Kreislauf-Forschung (DZHK) (A.B.)

European Research Council (ERC) Starting Grant PROTEOFIT (A.B.)

Deutsche Forschungsgemeinschaft (DFG) SFB1123-B5 (D.S.)

Bundesministerium für Bildung und Forschung (BMBF) (D.S.)

Free State of Bavaria (LMU Excellence strategy) (D.S.)

German Centre for Cardiovascular Research (DZHK) grants 81Z0600203 and 81X2600269 (D.S.)

European Research Council (ERC) under the European Union”s Horizon 2020 research and innovation program # 949017 (M.R.)

Helmholtz Association - Initiative and Networking Fund (M.R.)

EFSD/Novo Nordisk Foundation Future Leaders Award (M.R.)

European Stroke Research Foundation (ESRF) (A.H., C.R.)

Joachim Herz Foundation (J.G.P.L)

## Author contributions

B.A.A., A.C. and O.T.B. conceived and designed the study, which T.A.V. helped to guide. B.A.A. and J.Y. performed imaging experiments. J.G.P.L. built the customized spectrometer setup, while B.A.A., J.Y., J.G.P.L. and I.M. performed spectroscopic measurements. A.H., J.H., D.S., C.W. and C.R. provided clinical arterial biopsies and histology, while C.J., H.J., A.R.F., M.R., R.K.P., T.A.V. and A.B. provided other biological samples. K.M. and A.C.S. engineered the 4T1-iRFP720 cell line. I.M. and B.A.A. designed the illustrations. B.A.A., A.B., A.C. and O.T.B. drafted the manuscript, to which all authors contributed. All authors approved the submitted version.

