## Supplementary Information for "Macroscopic label-free biomedical imaging with shortwave infrared Raman scattering"

### Supporting Information
for

### Macroscopic label-free biomedical imaging with shortwave infrared Raman scattering

Bernardo A. Arús [1-5,*], Joycelyn Yiu [5], Jakob G. P. Lingg [5], Anja Hofmann [6], Amy R. Fumo [7,8], Honglei Ji [7,8], Carolin Jethwa [9], Roy J. Park [10], James Henderson [9,11], Kanuj Mishra [12], Iuliia Mukha[1-4,13], Andre C. Stiel [12,14], Donato Santovito [9,11], Christian Weber [9,11], Christian Reeps [6], Maria Rohm [7,8], Alexander Bartelt [7-9,11], Tulio A. Valdez [10], Andriy Chmyrov [1-5], Oliver T. Bruns [1-5,*]

[1] Department of Functional Imaging in Surgical Oncology, National Center for Tumor Diseases (NCT/UCC), Dresden, Germany

[2] German Cancer Research Center (DKFZ), Heidelberg, Germany

[3] Medizinische Fakultät and University Hospital Carl Gustav Carus, Technische Universität Dresden, Dresden, Germany

[4] Helmholtz Zentrum Dresden-Rossendorf (HZDR), Dresden, Germany

[5] Helmholtz Pioneer Campus, Helmholtz Zentrum München, Neuherberg, Germany

[6] Division of Vascular and Endovascular Surgery, Department of Visceral, Thoracic and Vascular Surgery, Faculty of Medicine and University Hospital Carl Gustav Carus, Technische Universität Dresden, Germany

[7] Institute for Diabetes and Cancer (IDC), Helmholtz Zentrum München, Neuherberg, Germany

[8] German Center for Diabetes Research (DZD), Neuherberg, Germany

[9] Institute for Cardiovascular Prevention (IPEK), Faculty of Medicine, Ludwig-Maximilians-Universität München (LMU), Munich, Germany

[10] Department of Otolaryngology, Head and Neck Surgery, Stanford University, Palo Alto, CA, USA

[11] German Center for Cardiovascular Research (DZHK), Partner Site Munich Heart Alliance, Munich, Germany [12] Institute of Biological and Medical Imaging, Helmholtz Zentrum München, Neuherberg, Germany

[13] Chuiko Institute of Surface Chemistry of the National Academy of Science of Ukraine, Kyiv, Ukraine

[14] Faculty of Biology and Pre-Clinical Medicine, University of Regensburg, Regensburg, Germany

*Bernardo A. Arús:
*Oliver T. Bruns:

#### Materials and Methods

Materials

Chemicals and laboratory consumables were purchased from either Sigma-Aldrich (Taufkirchen, Germany), Merck (Darmstadt, Germany), Carl Roth (Karlsruhe, Germany), Thermo Fisher Scientific (Roskilde, Denmark). Optical and optomechanical components were purchased from Edmund Optics (York, United Kingdom), Thorlabs (Bergkirchen, Germany), Qioptic (Göttingen, Germany), Lumics (Berlin, Germany), Schneider-Kreuznach (Bad Kreuznach, Germany), NKT Photonics (Birkerød, Denmark), or Teledyne Princeton Instruments (Trenton, NJ, USA). Mouse chow was purchased from Research Diets (New Brunswick, NJ, USA) or Altromin (Lage, Germany).

System for wide-field macroscopic imaging based on SWIR Raman scattering

We developed the system using a design similar to that of systems to detect SWIR fluorescence, which we adapted for detection of Raman scattering by optimizing the wavelengths of detection corresponding to Raman bands upon specific illumination wavelengths, and by using a low-noise, InGaAs-based detector capable of detecting SWIR radiation during long exposure times.

Samples were illuminated using the following lasers (Lumics) to obtain the wavelengths indicated in the figure legends: 672 nm, LU0680D010-E10FH; 785 nm, LU0785D250-U70AN; 892 nm, LU0890D400-U10AF; 938 nm, LU0940D250-U70AF; 968 nm, LU0980D350-U30AF; and 1064 nm, LU1064D350-U70AF. Lasers were connected to a 7 × 1 fan-out fiber-optic bundle (catalog no. BF76LS01, Thorlabs) with a core diameter of 600 μm. The laser light traveled from the common end of the multi-fiber bundle into an excitation cube (catalog no. KCB1EC/M, Thorlabs), it was reflected off a mirror (catalog no. BBE1-E03, Thorlabs), then it passed through a positive achromat (catalog no. AC254-050-B, Thorlabs). The wavelength of the laser output was cleaned up using appropriate short-pass filters to prevent detection of the laser light, such as a 900-nm filter for illumination at 892 nm. From the achromat, the laser light passed through an engineered diffuser (catalog no. ED1-S20-MD, Thorlabs) to provide uniform illumination on the field of view.

Depending on the experiment, the maximum laser power density varied roughly between 50 and 150 mW/cm$^2$, as measured using a PM100D power meter equipped with a S130C sensor (Thorlabs). The densities remained within international exposure limits (International Commission on Non-Ionizing Radiation 2013).

The nitrogen-cooled camera NIRvana LN (Princeton Instruments) was mounted on a rail carriage (catalog no. G026428000, Qioptic) parallel to an optical table (catalog nos. SDR7590 and B7590L, Thorlabs). This camera was chosen because of its deep cooling, low dark current and low read noise, which provide high signal-to-noise ratios at long exposures in a single frame. We verified that the simpler, air-cooled portable NIRvana HS (Princeton Instruments) can also detect Raman scattering (**Fig. S28**). An f/2.8, 50-mm lens capable of transmitting SWIR radiation (SWIRON 2.8/50, Schneider-Kreuznach) was attached to the camera using an F-mount adapter, while three 30-mm cage plates with removable holders (catalog no. CFH2/M, Thorlabs) for 1-inch optical filters (filter slots 1-3) were positioned in front of the camera lens to facilitate filter changes. A silver mirror in a right-angle kinematic elliptical holder (catalog nos. KCB1EC/M and PFE10-P01, Thorlabs) was mounted on the cage plate farthest from the lens in order to direct the laser light towards the optical table. A 60-mm cage plate with a removable holder for 2-inch optical filters (catalog no. LCFH2/M, Thorlabs) was attached to the side of the kinematic elliptical holder closer to the sample. Of the four filter positions, the one closest to the camera lens and the one closest to the sample (slots 1 and 4, respectively) were usually occupied by long-pass filters, which eliminated stray light from the laser; for example, the filters were long-pass 1000 nm when samples were illuminated at 968 nm. The two central filter positions were usually occupied by band-pass filters to define the detection band. With this set-up, we obtained fields of view typically around 56 cm$^2$, however a different choice of optics could enable larger or smaller fields of view, reflecting the scale-spanning character of wide-field imaging.

All optical components were covered with black aluminum foil (catalog no. BKF12, Thorlabs), and the system was enclosed within black nylon (catalog no. BK5, Thorlabs). All imaging was performed in a dark room.

Data acquisition

After detector temperature had stabilized at -190 ºC, data were acquired using Lightfield 6.15.1.2112 (Teledyne Princeton Instruments). Raman scattering in different bands was imaged sequentially with manual filter changes. Specific laser and filter specifications for each imaging experiment can be found in Table S1-S2. An illustrative data acquisition workflow for SWIR Raman imaging upon 892 nm consisted of the following: the laser was turned on, a first frame was acquired using the band-pass filter combination 1150-25 (center wavelength – width) to detect the Raman-silent region (**Fig. 1a**), the laser was turned off and the filter was changed, then the laser was turned back on and a second frame was acquired with the filter combination 1200-25 to detect the CH region. This process was repeated to acquire a third frame with the filter combination 1300-25 to detect the OH band. For data processing, dark frames were captured for each exposure time and flatfield images were captured for each laser configuration.

Data processing

Data were post-processed using Python 3 and the Fiji distribution of ImageJ 1.54f (US National Institutes of Health, Bethesda, MD, USA). First, dark and flatfield frames were generated by averaging 5-10 frames. Then, all image takes had their corresponding dark frames subtracted, after which they were further corrected for exposure time and divided by the power density map, which was obtained by multiplying together the flatfield frame and the maximum power density in the field of view. These steps yielded data in the form of counts/s/(mW/cm$^2$). Given the three-dimensional nature of *in vivo* wide-field imaging, images from *in vivo* experiments did not undergo flatfield correction, and were therefore represented in counts/s.

Image quantification was performed by drawing regions of interest including the signal of each sample, and the mean value or sum of intensities in each ROI was measured.

SWIR Raman imaging of isolated compounds

Olive oil (catalog no. 8873.1, Carl Roth) and water (0.75 mL of each) were transferred into a 1.5-mL microcentrifuge tube, and imaged in the wide-field SWIR Raman imaging setup upon 892 nm illumination. Powders from the isolated compounds, palmitic acid (catalog no. P0500, Sigma-Aldrich), cholesterol (catalog no. C8667, Sigma-Aldrich), hydroxyapatite (catalog no. 289396, Sigma-Aldrich), collagen from calf skin (catalog no. C9791, Sigma-Aldrich), and elastin from bovine neck ligament (catalog no. E1625, Sigma-Aldrich) were placed in a black 96-well plate (catalog no. 237108, Thermo Fisher Scientific), and imaged in the same setup at 1064 nm illumination.

Raman spectroscopy of isolated compounds

The isolated compounds were added to a black quartz cuvette (catalog no. Z801607, Merck), and spectra were acquired using the liquid nitrogen-cooled InGaAs line detector Pylon-IR 1024-1.7 (Teledyne Princeton Instruments) connected to a Czerny-Turner spectrograph (catalog no. HRS-300, Teledyne Princeton Instruments). A super-continuum white light laser (SuperK-Extreme, NKT Photonics) filtered with a tunable filter (SR-Extended-8HP LLTF Contrast, NKT Photonics) was used as excitation source. Spectra were acquired with illumination at 892 or 1064 nm, then cleaned up with, respectively, a 900-nm short-pass filter (catalog no. FESH0900, Thorlabs) or the combination of a band-pass filter 1064-10 nm (catalog no. FLH1064-10, Thorlabs) and 1100-nm short-pass filter (catalog no. 64-339, Edmund Optics). The photoemission was collected using a pair of silver-coated off-axis parabolic mirrors (catalog nos. MPD269-P01 and MPD269-P01-H, Thorlabs), which focus the light onto a 400-μm multimode fiber (catalog no. M28L01, Thorlabs) connected to the spectrograph. Either a 950-nm or two 1100-nm long-pass filters (catalog nos. FELH0950 and FELH1100, Thorlabs) were positioned in front of the fiber to further block the excitation light. Intensity correction takes were acquired with a calibration lamp (catalog no. SLS201L/M, Thorlabs). Background takes were also acquired. Background and intensity corrections were performed using Python3.

Animal procedures

Animal procedures were performed in conformity with institutional and national regulations and after approval by the government of Upper Bavaria. Experiments were carried out using four mouse strains: C57BL/6J, C57BL/6N, athymic nude, and Ob/Ob (all purchased from Charles River Laboratories, Germany). Animals were housed at a temperature of 23 °C and relative humidity of 45–65% on a 12/12 h light/dark cycle. Animals were imaged either after anesthesia with isoflurane at 4% in oxygen for induction and 1.5-2% for maintenance, or after euthanasia. During imaging of live animals, the mouse temperature was maintained between 35 and 37 ºC. Organ dissection was performed after animal euthanasia.

*Ex vivo* imaging of whole mice and excised organs

Male or female C57BL/6J mice were euthanized at 6 weeks old, had their hair removed, and were imaged in different positions using illumination at 892 nm, after which numerous tissues and organs were excised and imaged *ex vivo*. A similar procedure was used to image male and female 6-week-old C57BL/6J mice and Ob/Ob mice and their excised livers. To image lymph nodes in adipose tissue, a male 6-week-old C57BL/6J mouse was euthanized and imaged using illumination at 938 nm.

The Raman contrast of lipids and water was compared to autofluorescence by euthanizing male and female 4-week-old C57BL/6J mice, then imaging them followed by imaging their excised organs after illumination at 785, 892, 938, 968, or 1064 nm. Specific filters used in combination with these lasers are listed in Table S1-S2.

*In vivo* imaging of changes in mouse adipose tissue induced by fasting and refeeding

Female 8-week-old athymic mice had *ad libitum* access to food, whereupon they were anesthetized as described above and imaged using illumination at 938 nm. Then animals were returned to their home cages and fasted for 24 h. The animals were imaged again, returned to their cages and given *ad libitum* access to food. At 24 and 48 h later, the animals were imaged.

Cell culture conditions

The cell line 4T1 was purchased from the American Type Culture Collection (ATCC) (catalog no. CRL-2539). The cells were cultured in RPMI 1640 medium (catalog no. 11875093, Thermo Fisher Scientific) supplemented with 10% fetal bovine serum (catalog no. F4135, Sigma-Aldrich), 100 units $ml^{-1}$ penicillin and 100 µg $ml^{-1}$ streptomycin, at 37 °C in an atmosphere of 20% $O_2$ and 5% $CO_2$.

Generation of 4T1-iRFP720 cell line

Cultured 4T1 cells underwent an antibiotic kill curve assay employing varying concentrations of Geneticin G418 antibiotic (catalog no. 2039.2, Carl Roth) to determine the optimal selection conditions (G418 concentration ranging from 0 to 1000 µg/ml). The concentration resulting in complete cell death within a week (200 µg/ml) was selected. Subsequently, the cells were transfected with pCDNA3.0-iRFP720, which contains a neomycin gene conferring resistance to G418. The iRFP720 construct, optimized for mammalian codons, was obtained from Thermo Fisher Scientific and subcloned into pCDNA3.0. Transfection was performed using the Neon transfection system (Thermo Fisher Scientific). Following transfection, the cells were exposed to G418 for 7 days to select for those successfully incorporating the cassette. Afterwards, the cells were cultivated, and single-cell sorting was performed by gating the top 10% of 4T1 cells exhibiting high fluorescent intensity using a FACS Aria III (BD Bioscience). Selected cells were expanded for 2 weeks. Subsequently, individual clones were analyzed for the highest fluorescent intensity using a fluorescence microscope, and the superior clone was selected for further expansion. Once expanded, the chosen clone was cryopreserved in 5% dimethyl sulfoxide complete RPMI freezing medium (catalog no. R8758, Sigma-Aldrich).

Imaging of tumor growth in mice

For the tumor cell implantation, $4 \times 10^5$ cultured 4T1 or 4T1-iRFP cells were resuspended in 50 µL phosphate-buffered saline, and injected subcutaneously onto the flank of 7-week-old female athymic nude mice. After implantation, the mice were anesthetized and imaged in the wide-field SWIR Raman imaging setup upon 892 nm illumination. For iRFP excitation, 672 nm was used, and its emission tail was detected in the same setup using an 1100-nm long-pass filter. Imaging was performed daily.

Whole-body imaging of mice fed a choline-deficient high fat diet

Four male 7-week old C57BL/6N mice were fed a diet deficient in methionine and choline (catalog no. A06071302, Research Diets) for 6 weeks, at the end of which the mice were euthanized and their corpses were frozen at -80 $^{o}$C until analysis. Four littermates were fed standard chow as a control (catalog no. 1314, Altromin). The corpses were thawed, had their hair removed, positioned in the SWIR Raman imaging setup, and imaged using 938 nm illumination.

Imaging of dissected fatty liver from mice

Male 8-week old C57BL/6N mice were fed either a diet deficient in methionine and choline (catalog no. A06071302, Research Diets) for 7 weeks or a diet containing 60% kcal% fat (catalog no. D12492, Research Diets) for 16 weeks. As controls, mice were fed standard chow. Six mice per group were utilized. Mice were euthanized at the end of the diet protocol, after which the ones from the group receiving a high-fat diet and the corresponding controls were perfused with phosphate-buffered saline. Livers were removed from all animals and a piece of tissue was excised, weighed, snap-frozen in liquid nitrogen, and stored at -

80 $^{o}$C until analysis. Livers were thawed on ice and imaged using illumination at 1064 nm. Images were flatfield-corrected. In the CH region images, we drew regions of interest covering the entire sample and, in order to substract background laser counts, four regions of interest covering areas surrounding the sample.

Porcine tissue collection

Porcine head and brain samples were obtained from a local slaughterhouse and kept at 4°C. They were transported to the laboratory the morning following slaughter. In the laboratory, the facial nerve was carefully exposed *in situ* and subsequently subjected to imaging. No tissue fixation was performed.

Imaging of atherosclerotic plaques in human arteries

Surgical biopsies from women 72-81 years old with high-grade stenosis who underwent carotid or iliofemoral endarterectomy were obtained with their informed consent and the approval of the ethics committee of the Technische Universität Dresden and Ludwig Maximilian Universität München. Biopsies were placed in ice-cold phosphate-buffered saline, excised at the largest site of the plaque, snap-frozen in liquid nitrogen and stored at -80 $^{o}$C until analysis. Samples were thawed and imaged using illumination at 1064 nm.

To assess the specificity of the calcification signal, surgical biopsies were imaged before and after 10% neutral buffered formalin fixation (catalog no. HT501128, Merck), followed by 14-day decalcification with Osteosoft (catalog no. 101728, Sigma-Aldrich).

Histological analysis

Mouse liver specimens were fixed in 4 % (w/v) neutrally buffered formalin, embedded in paraffin, and cut into 3 µm sections. Human atherosclerotic lesions were fixed in 4 % (w/v) phosphate-buffered paraformaldehyde, decalcified, embedded in paraffin, and cut into 5 µm sections. Sections from both types of tissues were stained with hematoxylin and eosin (HE), using a HistoCore SPECTRA ST automated slide stainer (Leica Biosystems) according to the manufacturer's instructions. Stained tissue sections were scanned with an AxioScan 7 digital slide scanner (Zeiss, Oberkochen, Germany) equipped with a 20x magnification objective.

Statistical analysis

Differences between groups were statistically analyzed using an unpaired, two-tailed t test. $P < 0.05$ was considered significant. All analyses were performed using GraphPad Prism v. 10.2.2. Data are represented as mean ± SD.

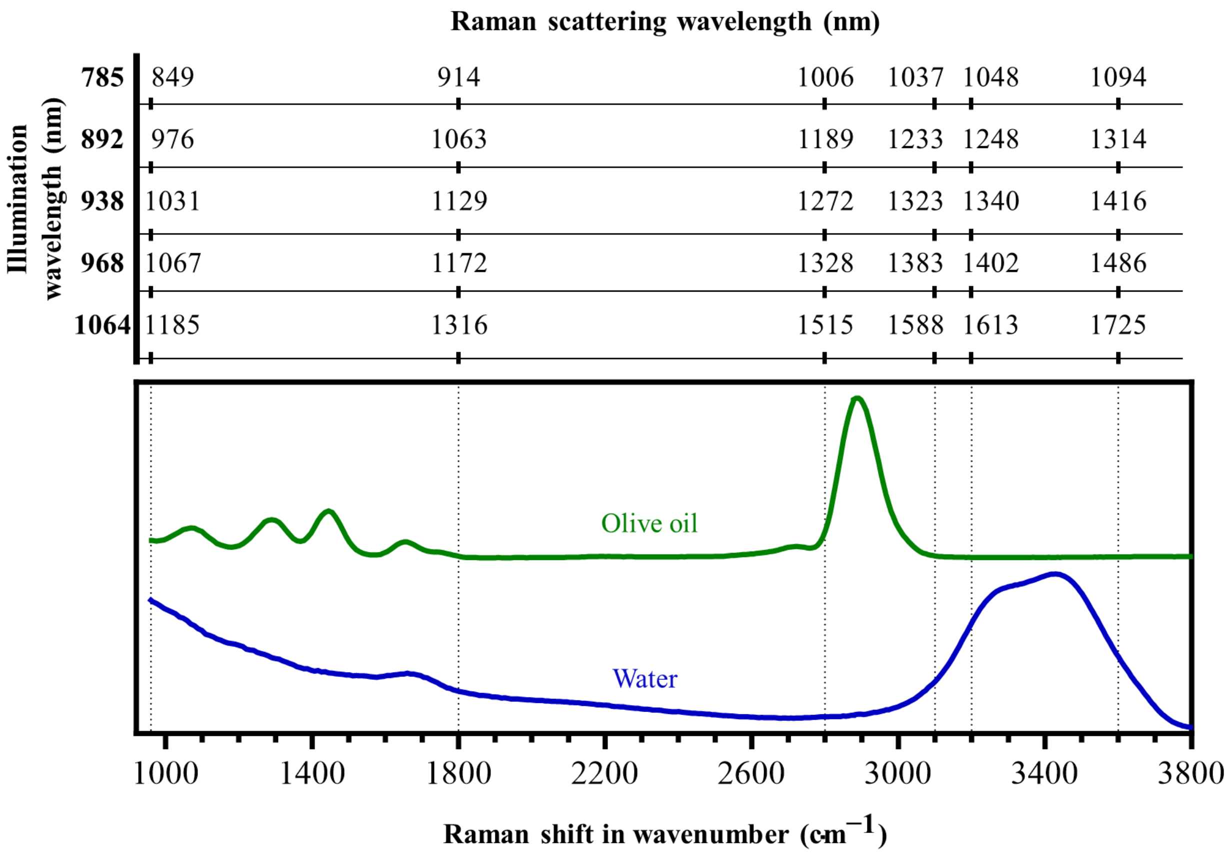

**Fig. S1**. Raman shift conversion from wavenumber to wavelength. Raman spectrum of olive oil and water acquired with 892 nm illumination and represented in wavenumbers in the lower (primary) x-axis. Following Equation 1, the primary x-axis was converted to the Raman wavelength shift for each illumination wavelength used in this work and displayed in the upper (secondary) x-axis. Arbitrary boundaries of the fingerprint (960 to 1800 cm$^{-1}$), silent (1800 to 2800 cm$^{-1}$), CH (2800 to 3100 cm$^{-1}$), and OH (3200 to 3600 cm$^{-1}$) Raman regions were selected (dashes vertical lines) to illustrate the shifts in wavelengths.

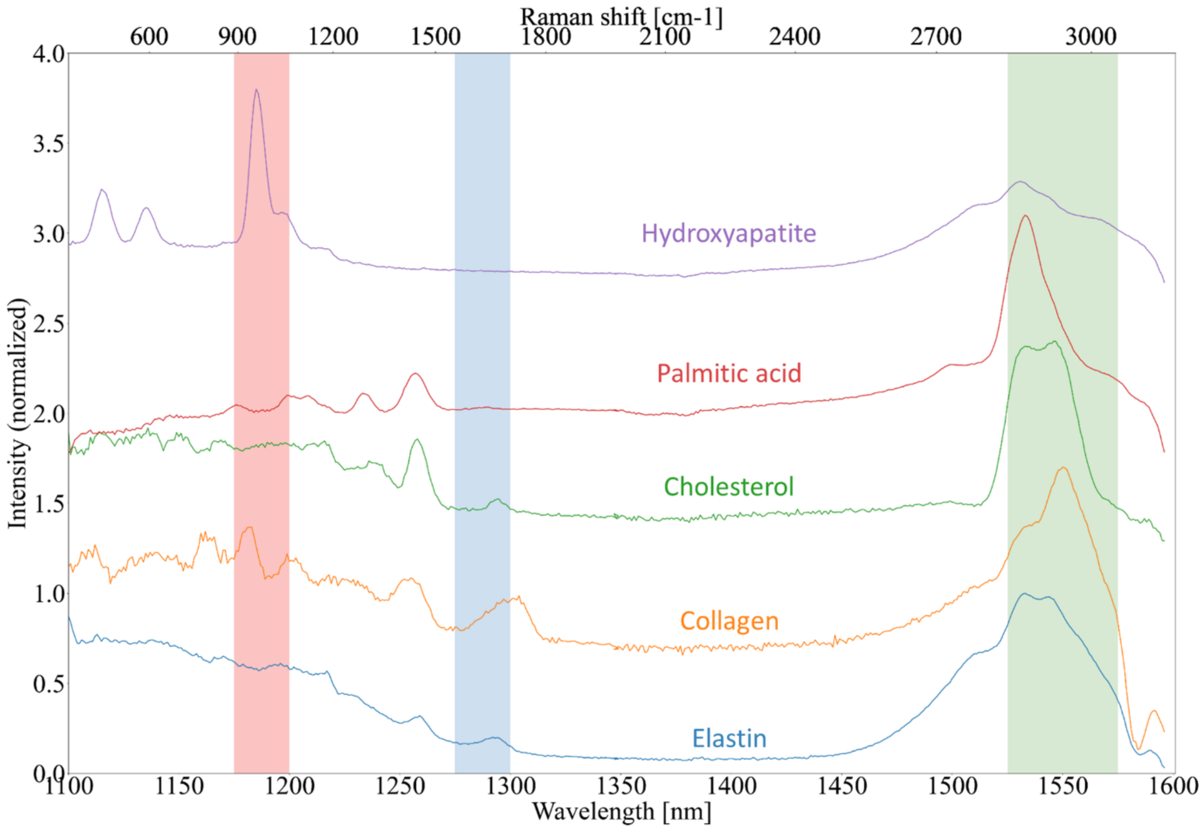

**Fig. S2**. Raman spectra of calcification-associated hydroxyapatite, the lipids palmitic acid and cholesterol, as well as the proteins collagen and elastin measured using illumination at 1064 nm. The primary (lower) x-axis displays the spectra in the detected wavelength regime, whereas the secondary (upper) x-axis represents the Raman shift upon 1064 nm illumination. The full-width at half-maximum of the band-pass filters used in Fig. S3 to detect the $PO_4^{-3}$, Amide I, and CH bands are labeled in red, green, and blue, respectively.

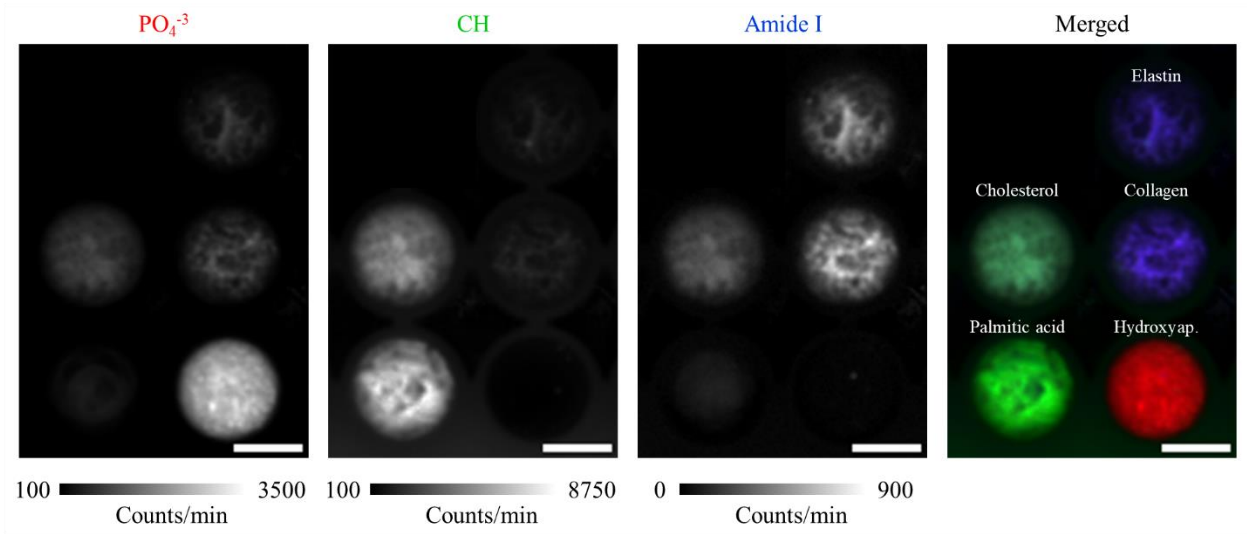

**Fig. S3**. SWIR Raman imaging of the compounds from Fig. S2. The powder compounds were placed into a polystyrene 96-well plate, and imaged in the SWIR Raman setup upon illumination at 1064 nm. Hydroxyap.: hydroxyapatite. Scale bar: 0.5 cm.

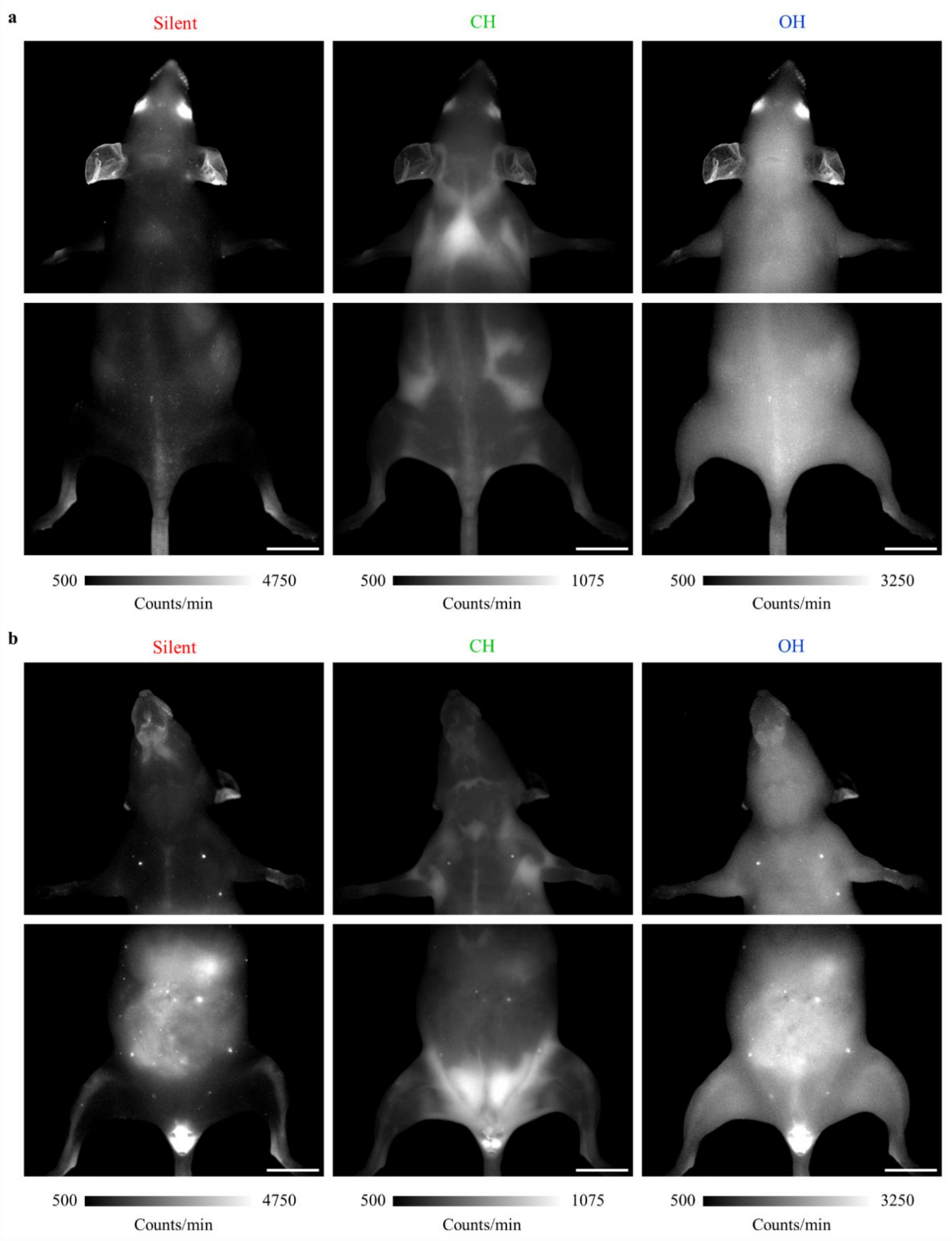

**Fig. S4**. Single-band images of the **(a)** dorsal and **(b)** ventral views of the mouse shown in Fig. 1 d. Scale bar: 1 cm.

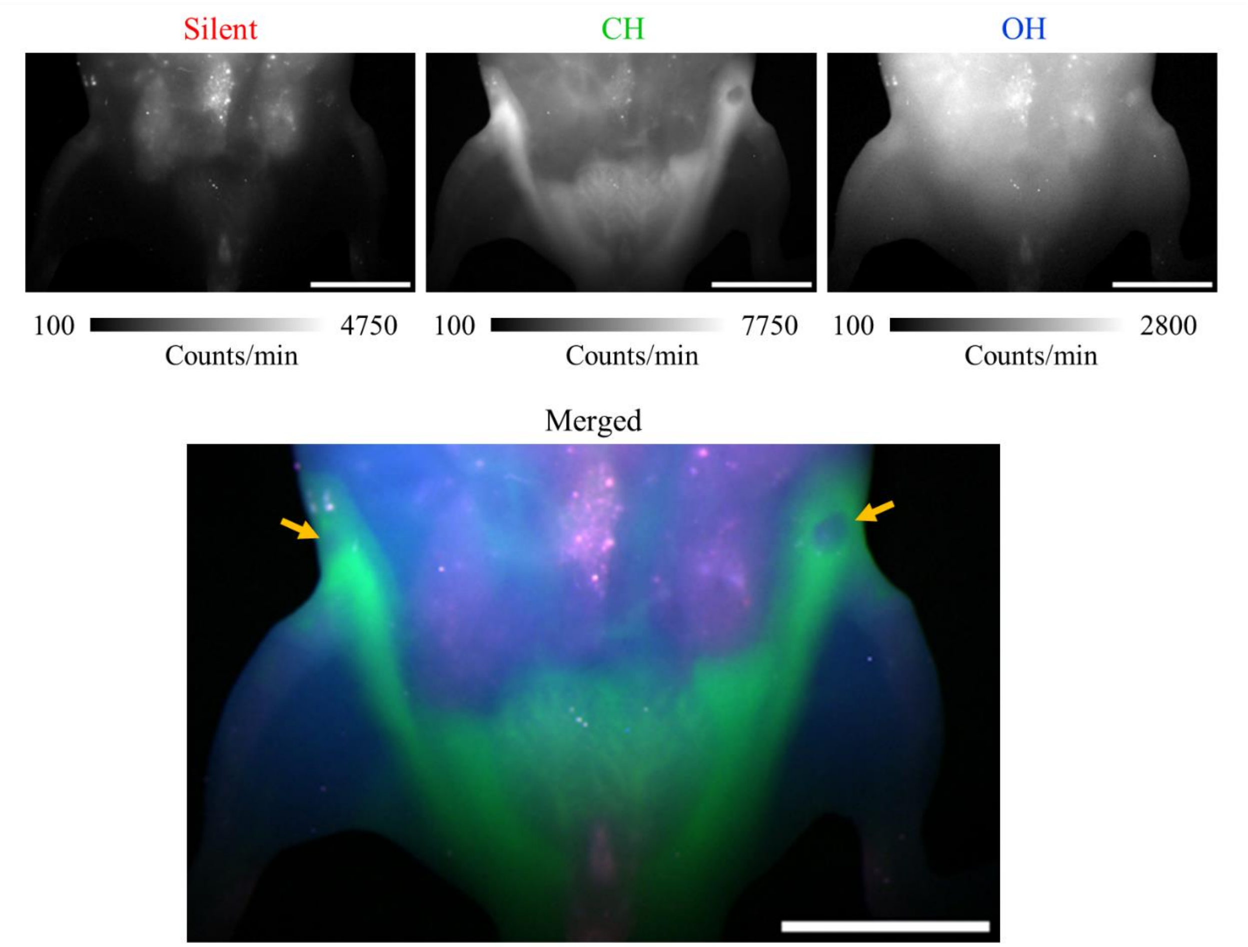

**Fig. S5.** Non-invasive inguinal lymph node (indicated by the yellow arrows) visualization in an intact C57BL/6J mouse based (ventral view) on SWIR Raman chemical contrast upon 938 nm illumination. Scale bar: 1 cm.

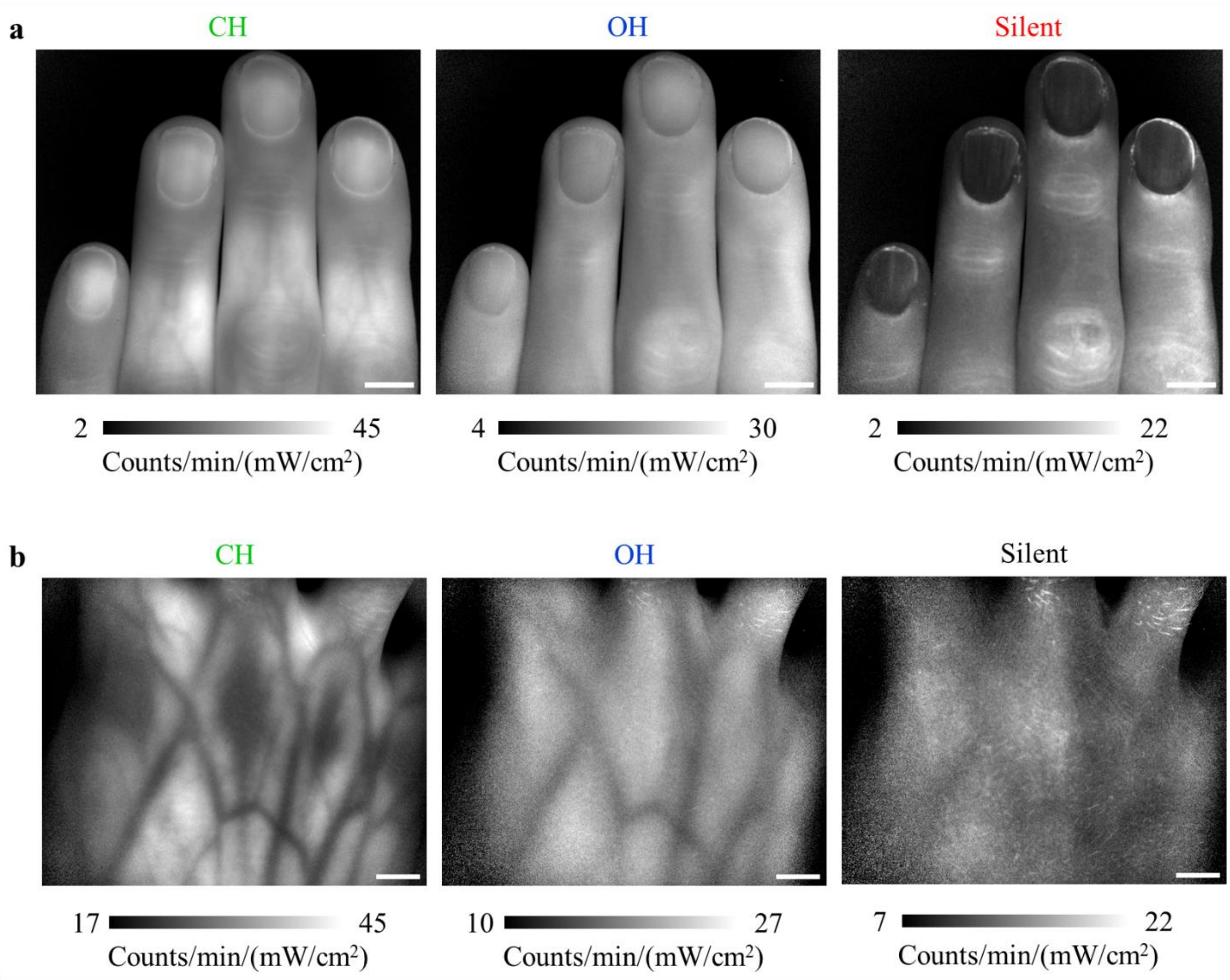

**Fig. S6**. Single-band images of the human hands shown in **(a)** Fig. 1 e and **(b)** Fig 1 f. The silent region of panel (b) was not shown in Fig 1 f because the autofluorescence mostly originated from the skin surface, thus impairing the visualization of the subcutaneous structures from the CH and OH Raman bands in the multicolor image. Scale bar: 1 cm.

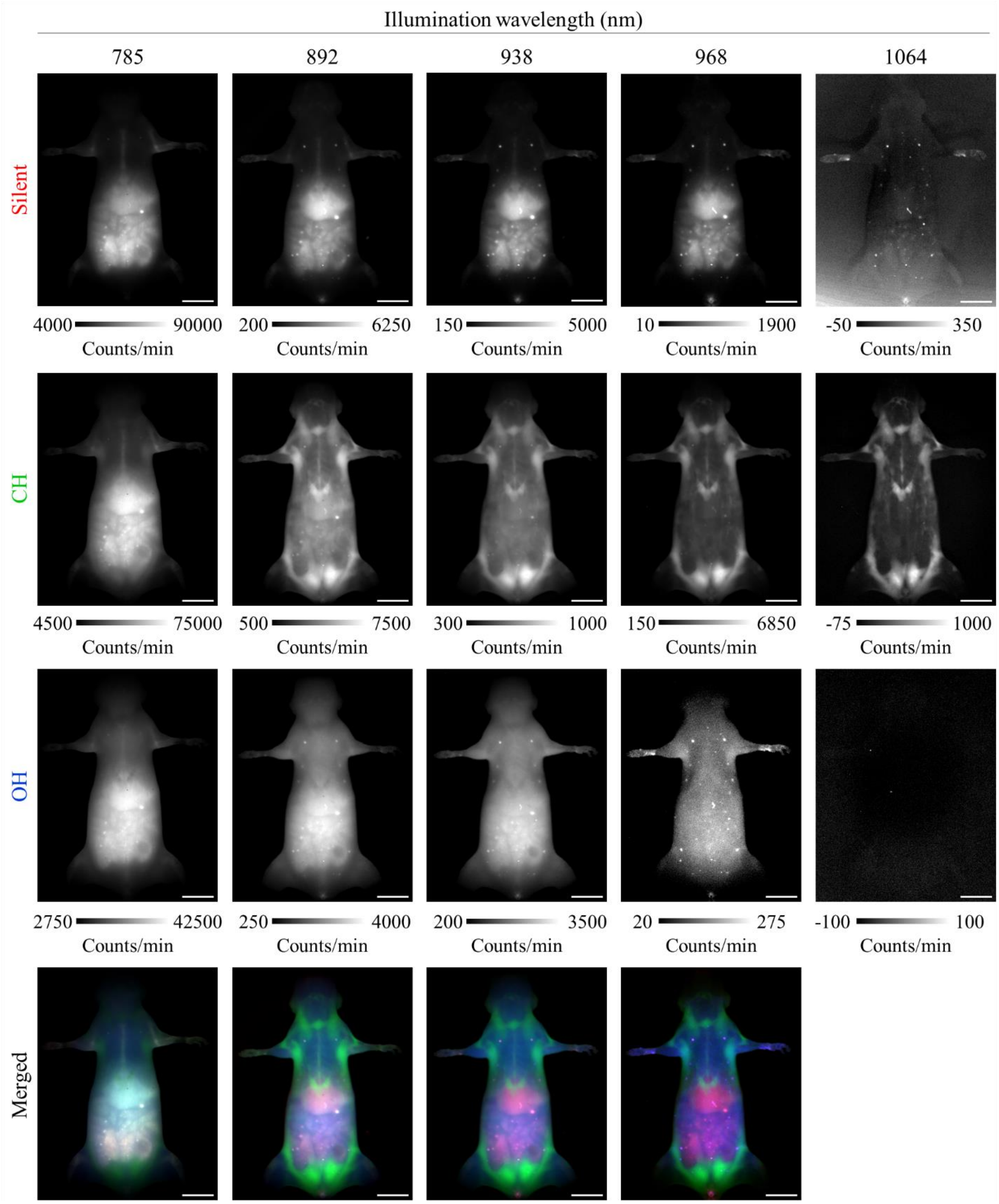

**Fig. S7**. Single-band images of the mouse images shown in Fig. 2 a-b (ventral view), with the addition of 1064 nm illumination, which was not shown in Fig. 2 a as a merged image because its OH region lies beyond the detector limit. The data are representative of 2 mice. Scale bar: 1 cm.

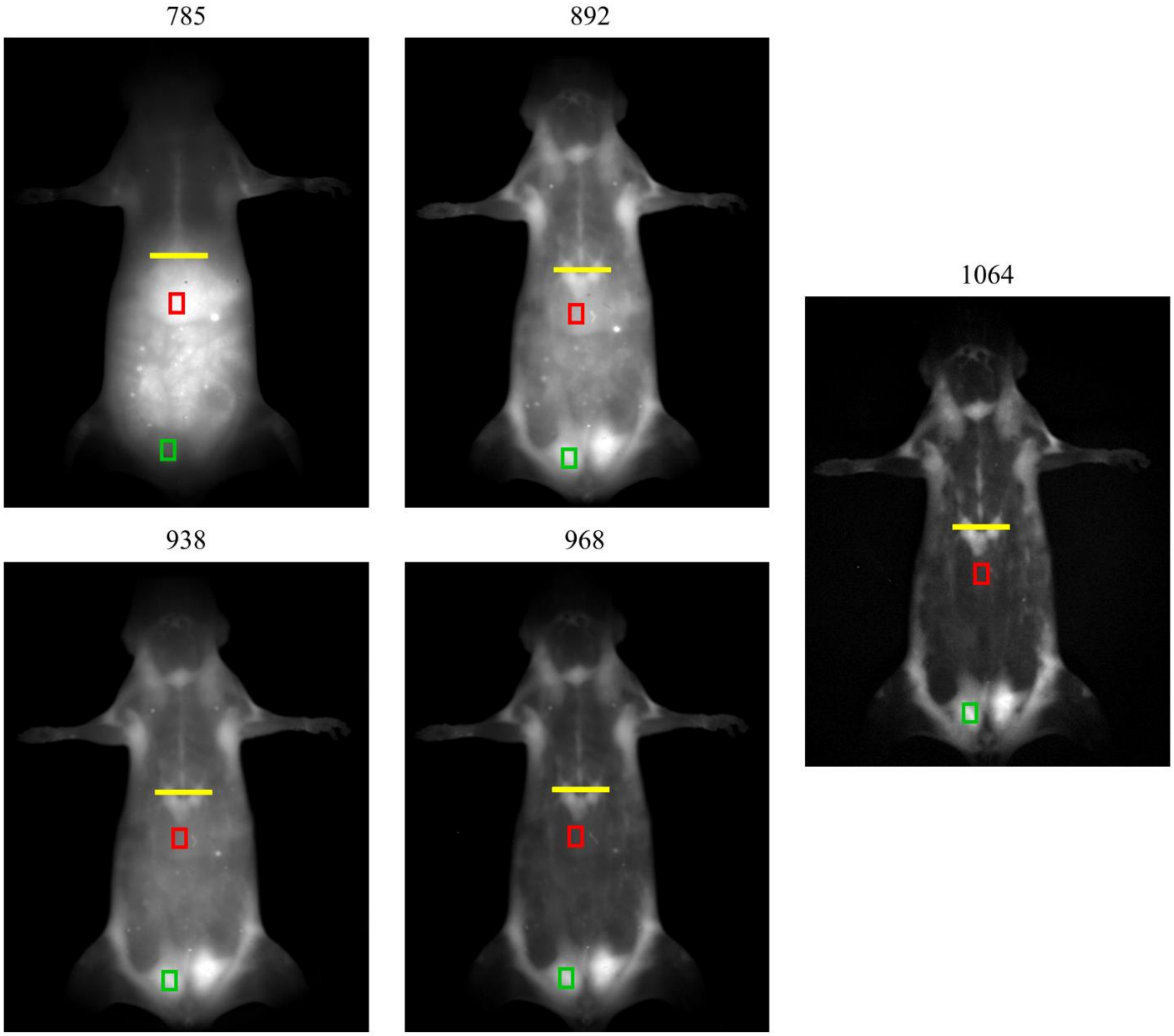

**Fig. S8**. Regions of interest (ROIs) used for data extraction for Fig. 2 c-d overlaid with the CH SWIR Raman images shown in Fig. 2 b and Fig. S6 (ventral view). The yellow line represents the ROI from which the line profiles shown in Fig. 2 c were extracted. In addition, the red and green rectangles respectively represent the ROIs from which the mean values of fat pad and liver were extracted for Fig. 2 d.

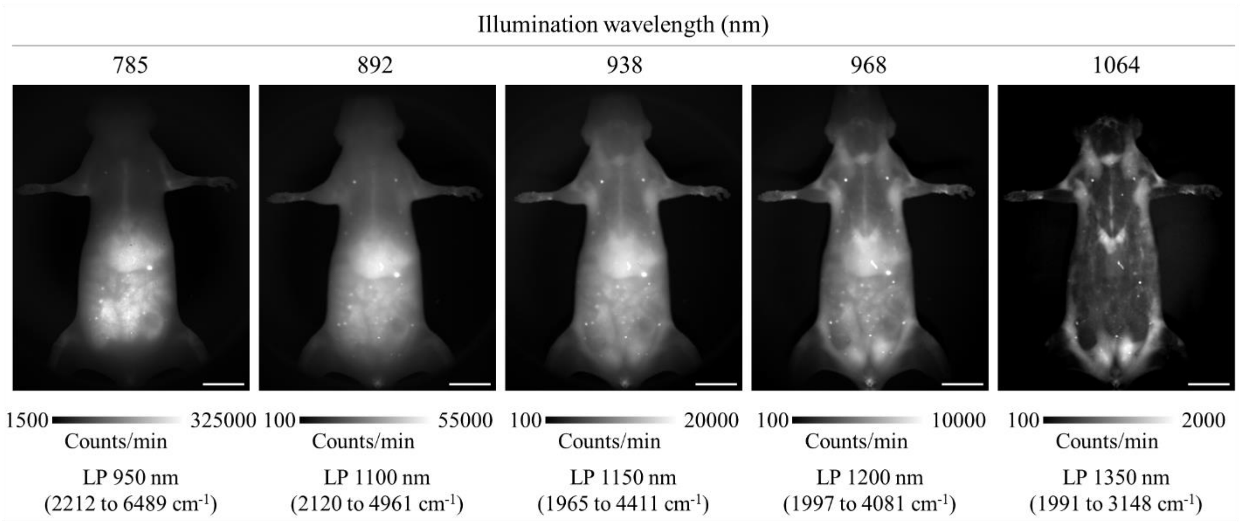

**Fig. S9.** Images acquired using long-pass (LP) filters, after illumination at the same wavelengths as in Fig. 2 a-b and Fig. S6 (ventral view). These LP filters cover wavelengths comprising the incident wavelength + Raman shift corresponding to wavenumbers above 1965 $cm^{-1}$ (the detection range, in wavenumbers, is shown in parenthesis, considering the detection limit of the camera as 1600 nm). As a consequence, these LP images detected the added signal from Raman-silent, CH and OH Raman bands for each illumination wavelength within a single frame. The OH band after illumination at 1064 nm lies beyond the detector's limit, so it does not contribute to the wide-band signal. The wavenumbers displayed here were rounded to the nearest integer, and can also be found in Table S2**.** The same mouse used in Fig. 2 a and Supplementary Fig. S7 was imaged. The data are representative of 2 mice. Scale bar: 1 cm.

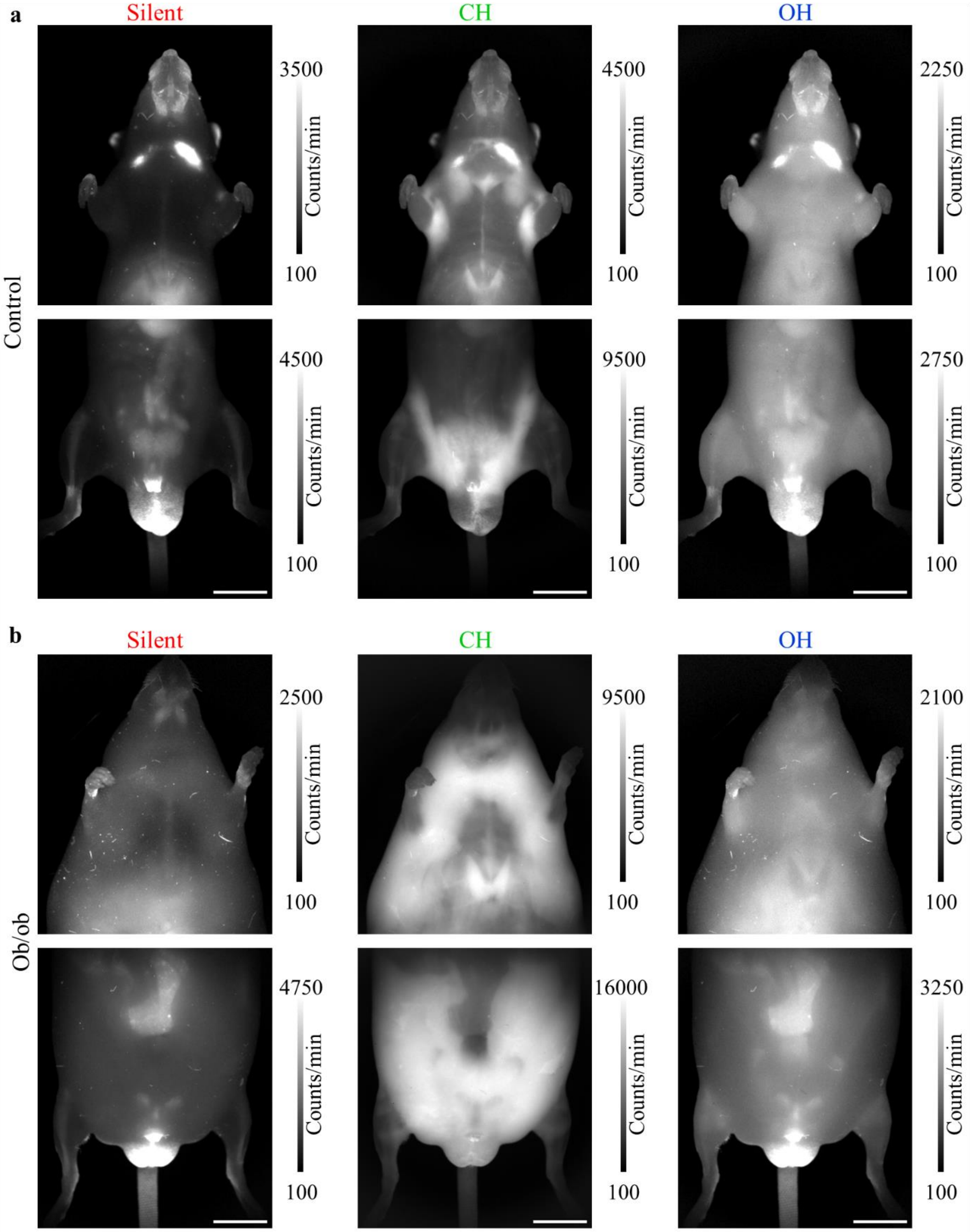

**Fig. S10**. Single-band images of the mice shown in Fig. 3 a (ventral view). Representative of 2 mice per group. Scale bar: 1 cm.

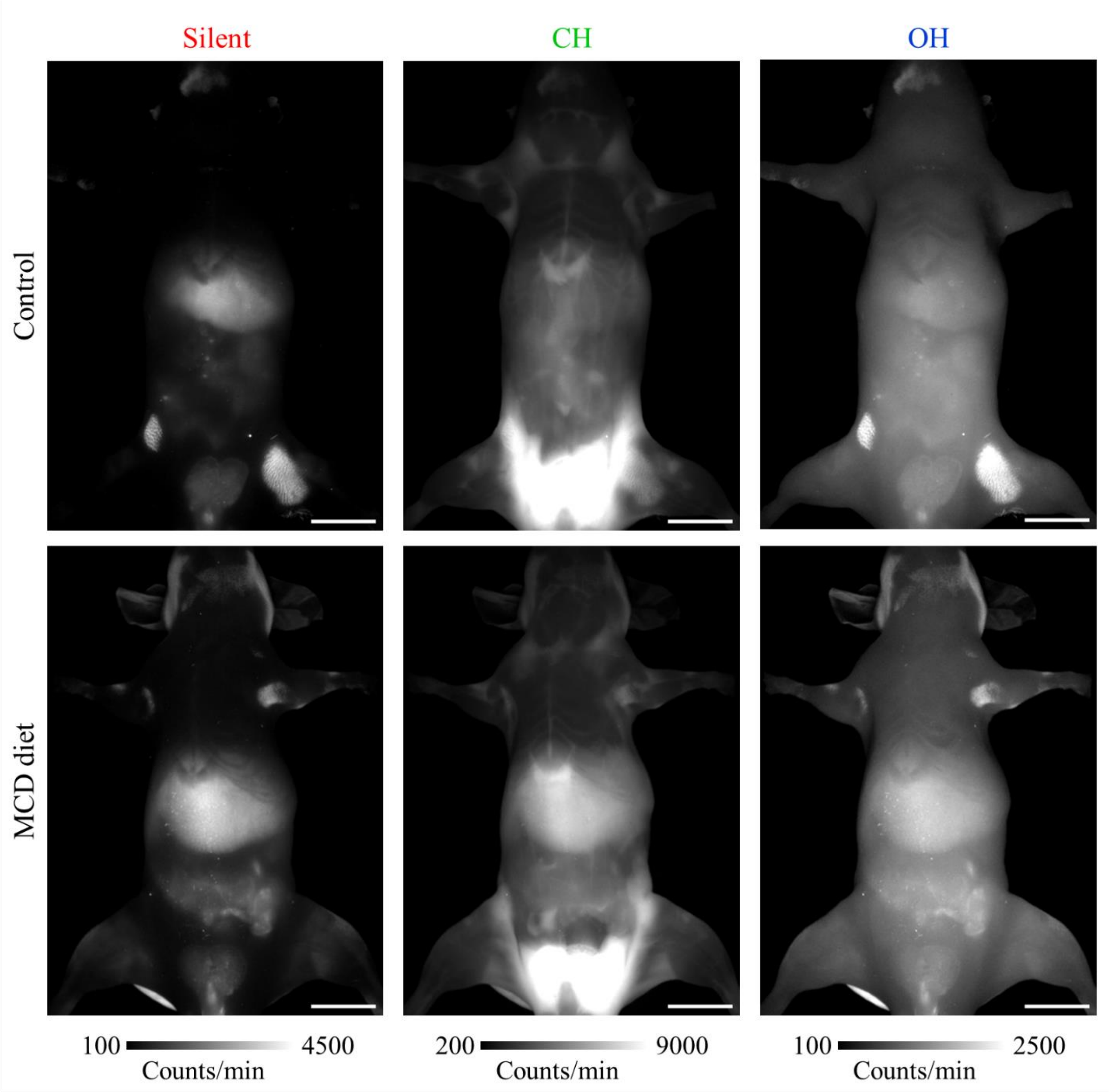

**Fig. S11.** Single-band images of the mice in Fig. 3 b-c (ventral view). The data are representative of 4 mice per group, and the other replicates are shown in Fig. S12**,** while the region of interest (ROI) selected for liver CH Raman intensity measurements (Fig. 3 d) are shown in Fig. S13. Scale bar: 1 cm.

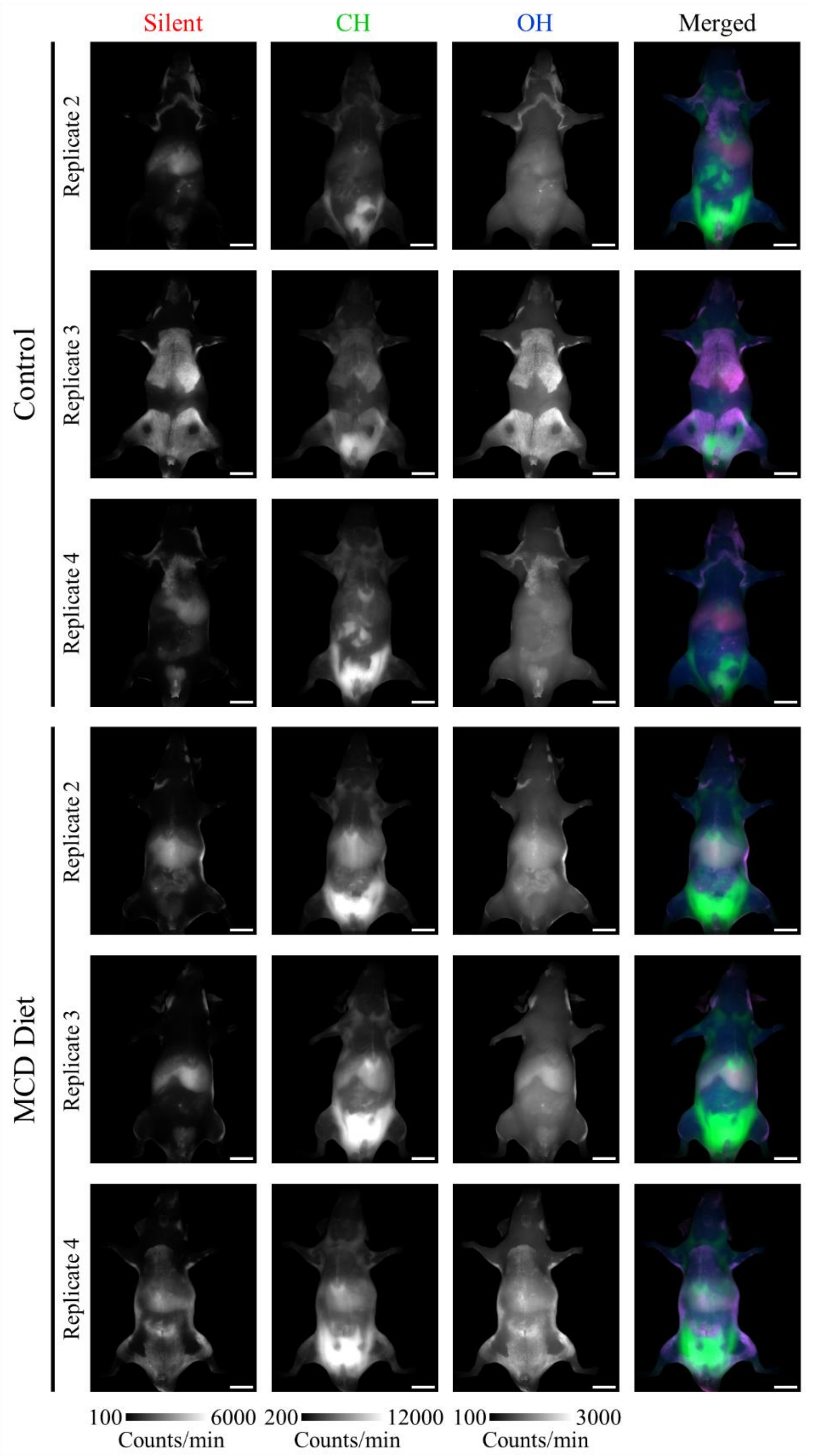

**Fig. S12.** Single-band and merged images of biological replicates of mice under a diet deficient in methionine and choline (MCD) or on a normal diet (control), corresponding to the experiment shown in Fig. 3 b-d and Fig. S11 (ventral view). Scale bar: 1 cm.

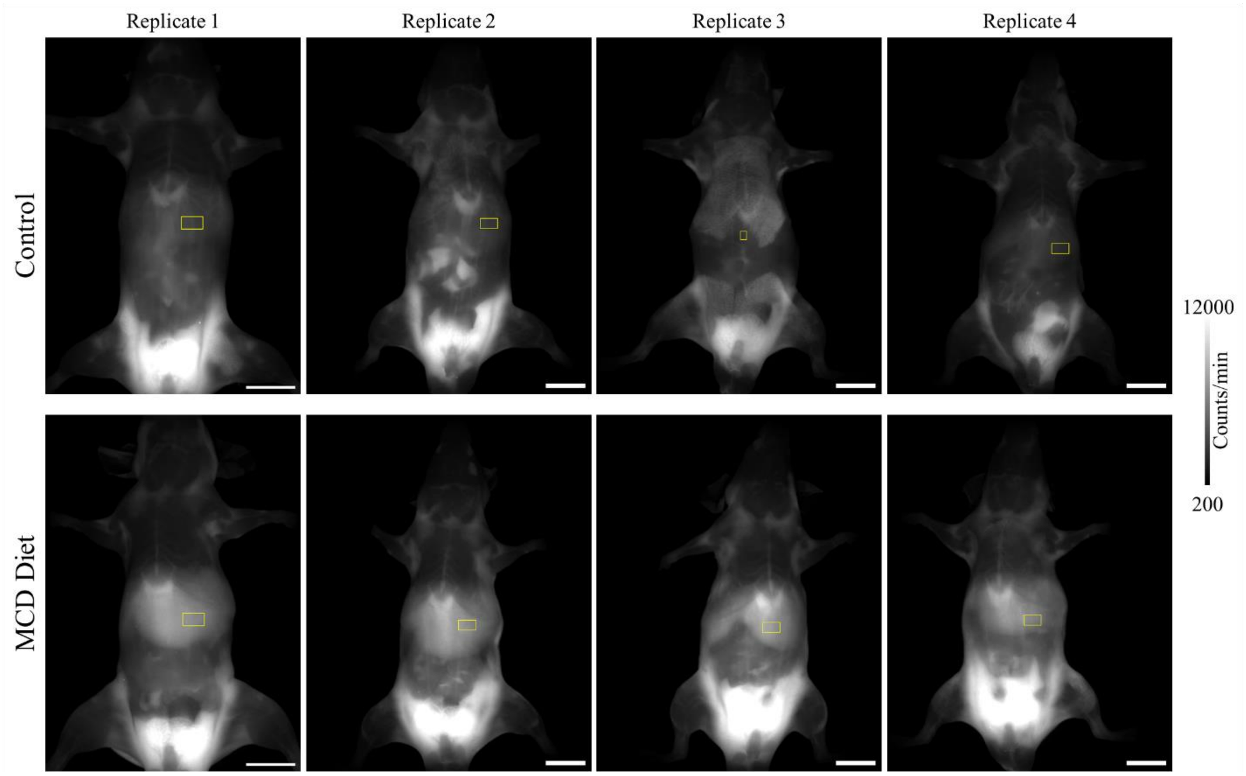

**Fig. S13.** Regions of interest (ROIs, yellow rectangles) used for data extraction for Fig. 3 d overlaid with the CH SWIR Raman images (ventral view) shown in Fig. 3 b-c and Fig. S12. Given the strong presence of melanin autofluorescence on the skin area covering the liver of replicate 3 (control), a small ROI was selected to avoid the interference of melanin onto the measured CH region intensity. To keep the ROI size consistent, similarly sized and positioned ROIs were selected for the other replicates. Scale bar: 1 cm.

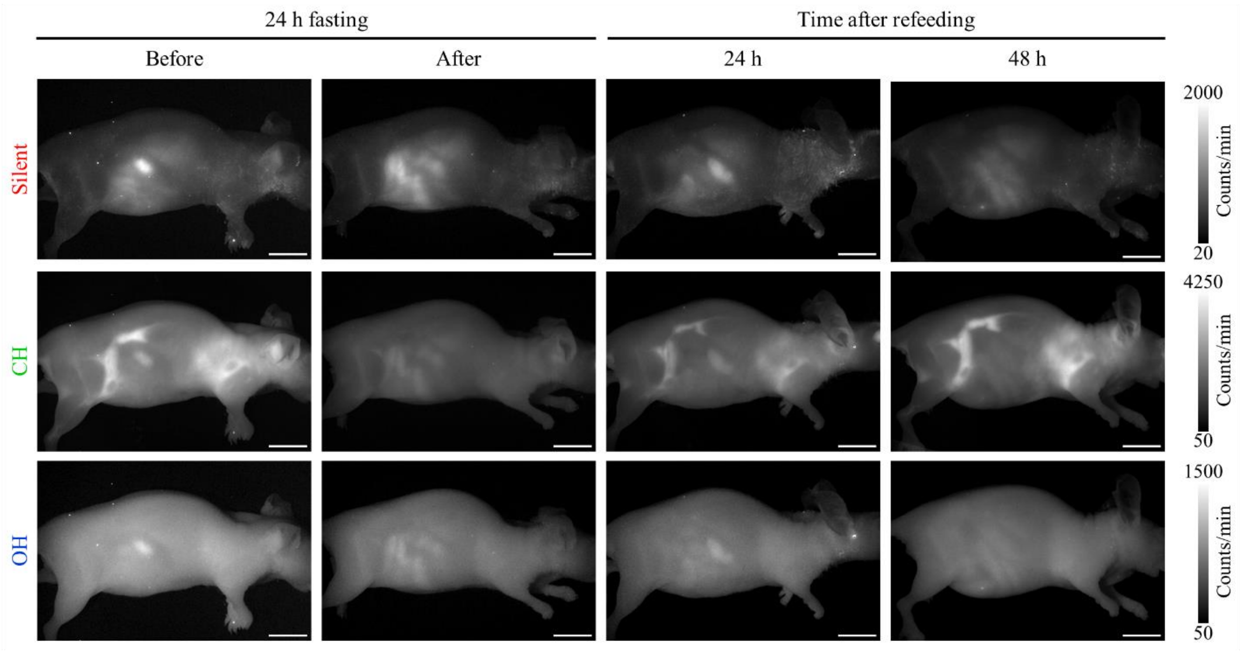

**Fig. S14.** Single-band images (right lateral view) of the anesthetized mouse images shown in Fig. 3 e. Scale bar: 1 cm.

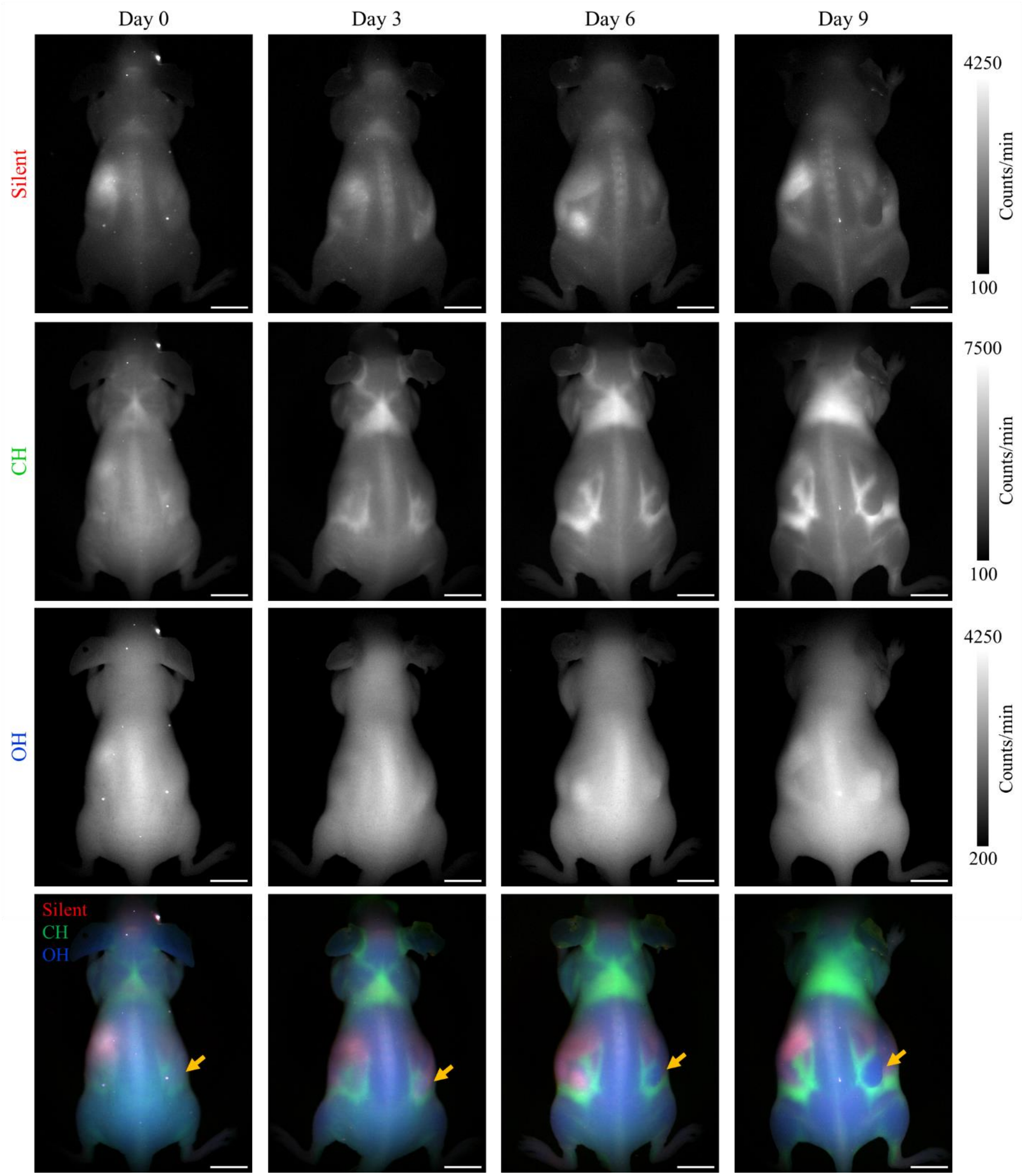

**Fig. S15.** SWIR Raman imaging of anesthetized mice bearing subcutaneous xenografts of 4T1 breast cancer cells in the right flank. Animals were imaged at the indicated number of days relative to tumor cell implantation. The arrow indicates the growing tumor. Results are representative of four animals. Scale bar: 1 cm.

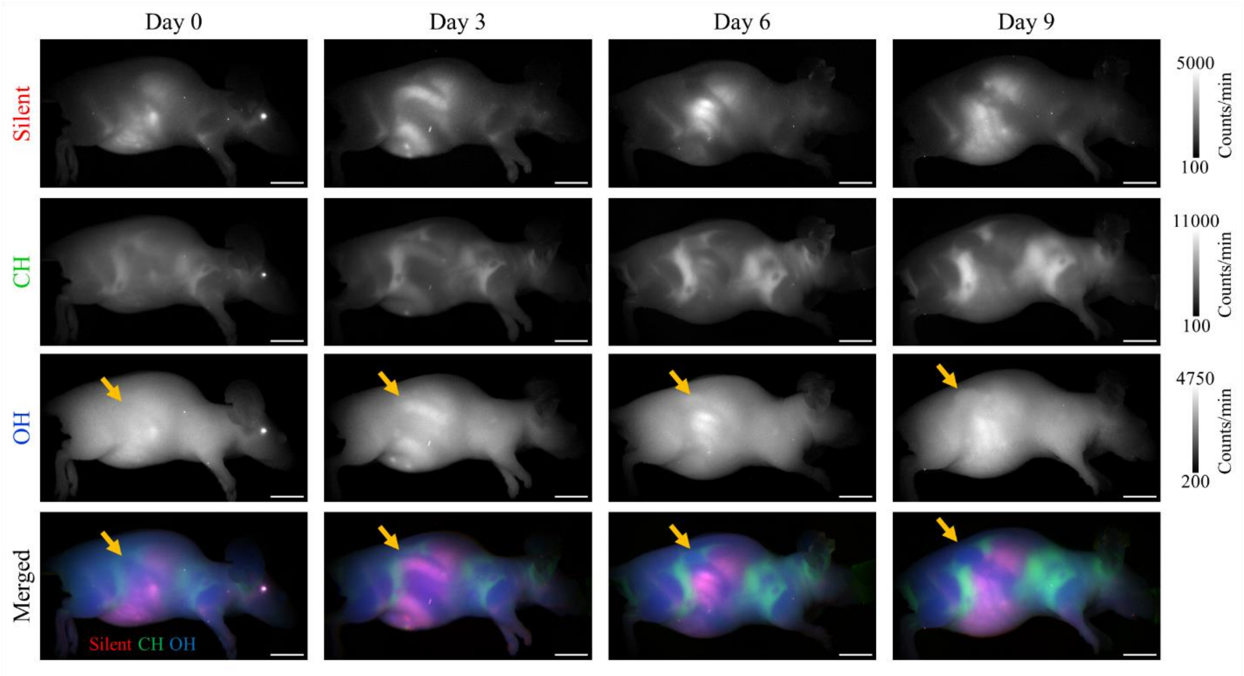

**Fig. S16.** Right lateral view of the mouse shown in Fig. S15. The arrow indicates the tumor location. Results are representative of four animals. Scale bar: 1 cm.

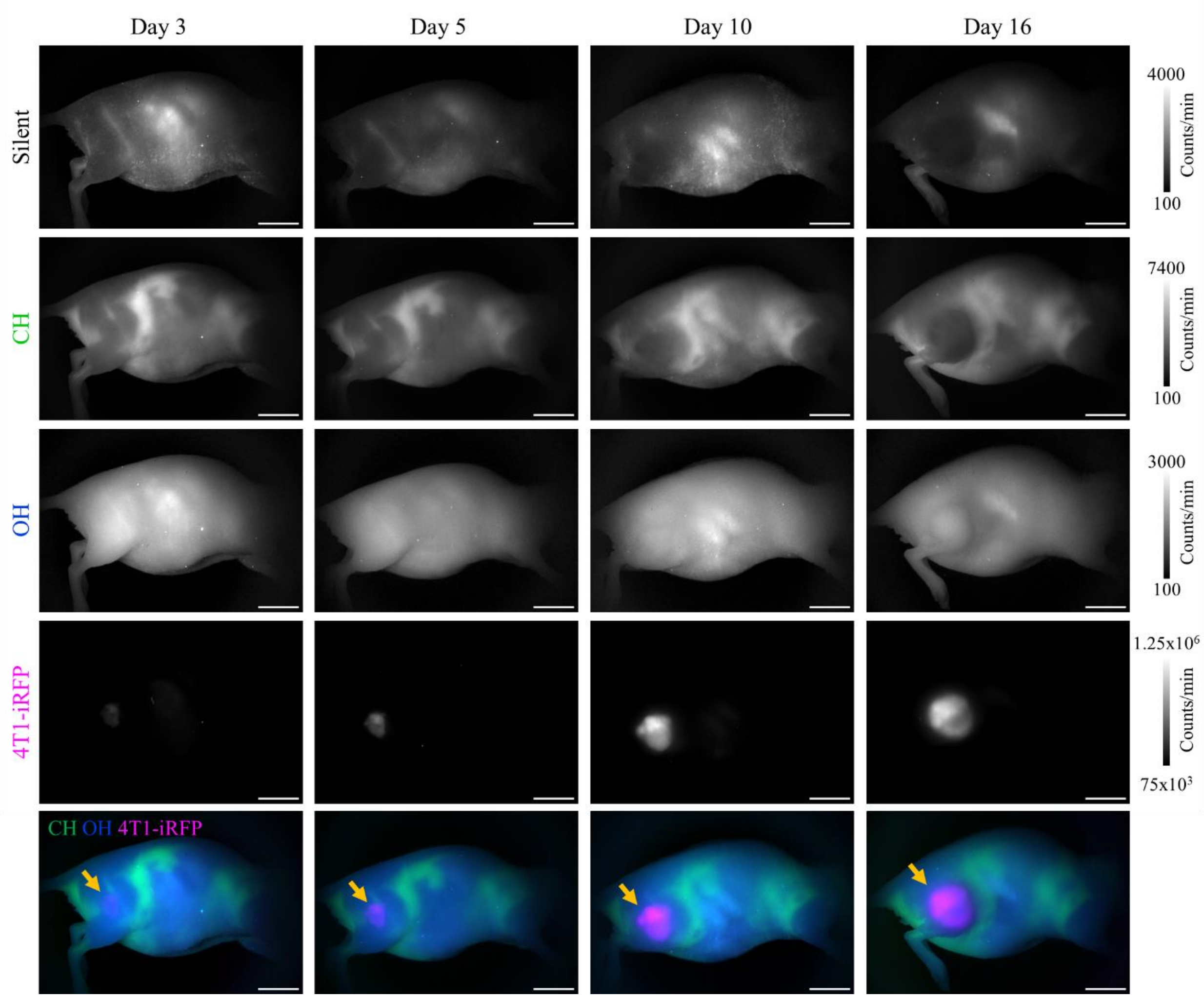

**Fig. S17**. Combination of wide-field Raman imaging and fluorescence imaging of mice like those in Fig S16, except that the 4T1 tumor cells were expressing the fluorescent protein iRFP720 and were injected onto the right leg. Results are representative of two animals. Configurations for Raman and fluorescence imaging are listed in Tables S1-2. Scale bar: 1 cm.

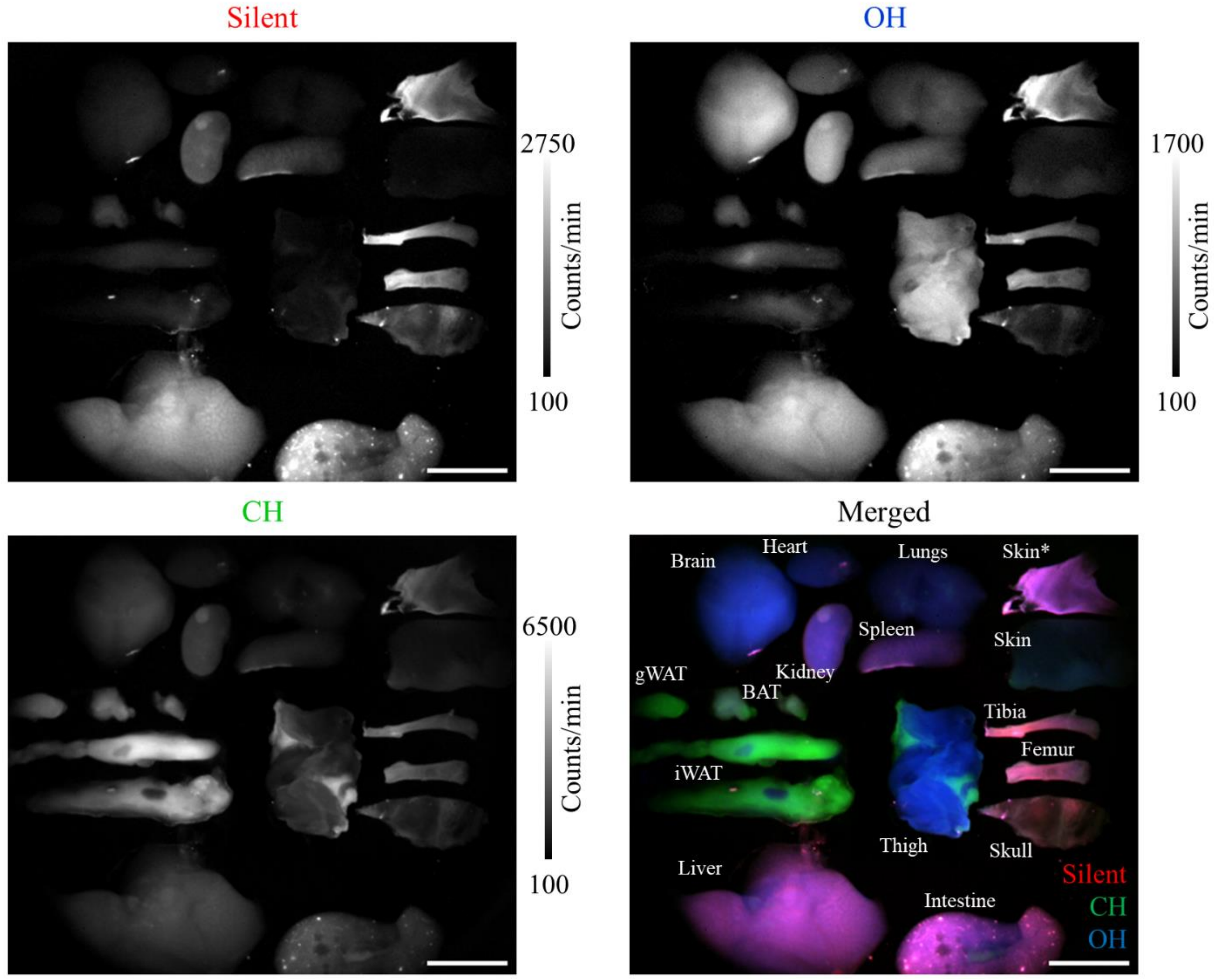

**Fig. S18.** Wide-field SWIR Raman imaging of organs from a healthy C57BL/6J mouse upon 892 nm illumination. Legend: BAT, brown adipose tissue; gWAT, gonadal white adipose tissue; iWAT, inguinal white adipose tissue; skin*, skin with melanin patch. The data are representative of 2 mice. Scale bar: 1 cm.

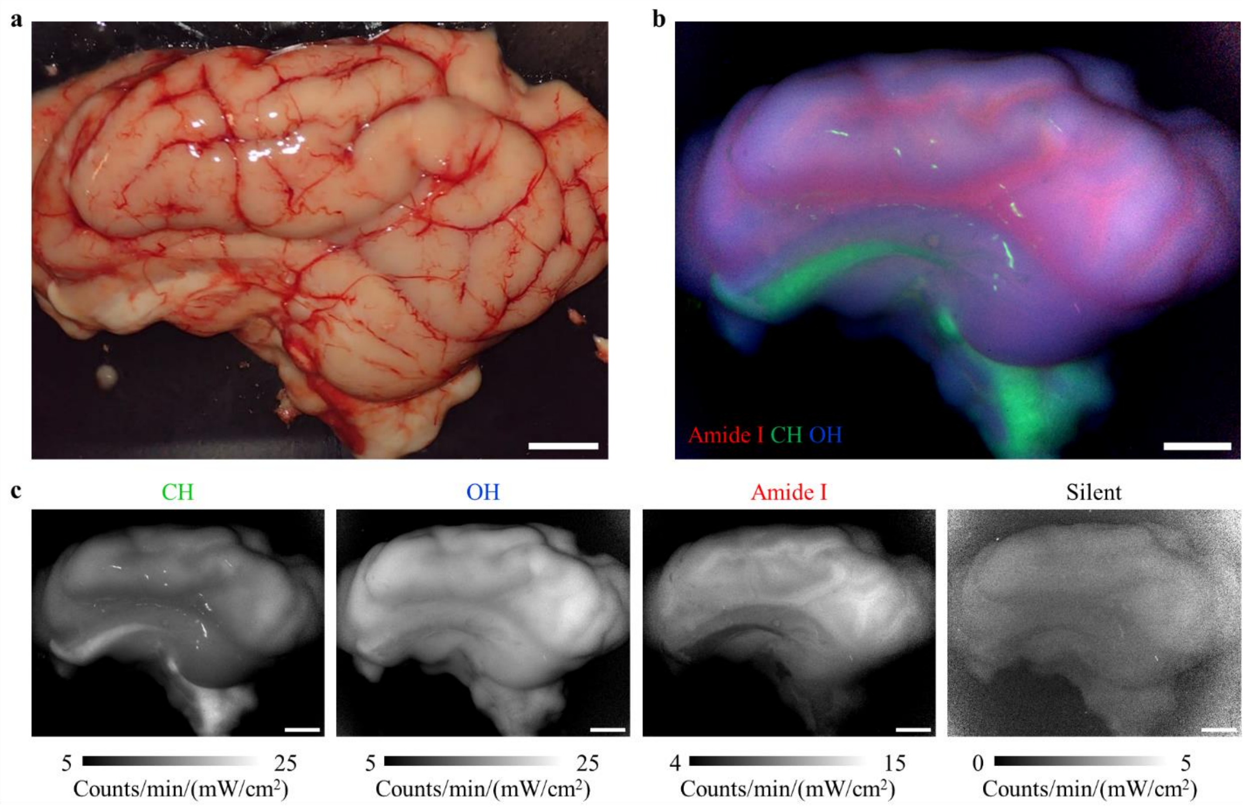

**Fig. S19.** Porcine brain outer cortex (lateral view of the left hemisphere) imaged **(a)** in the visible and **(b-c)** using SWIR Raman imaging. Scale bar: 1 cm.

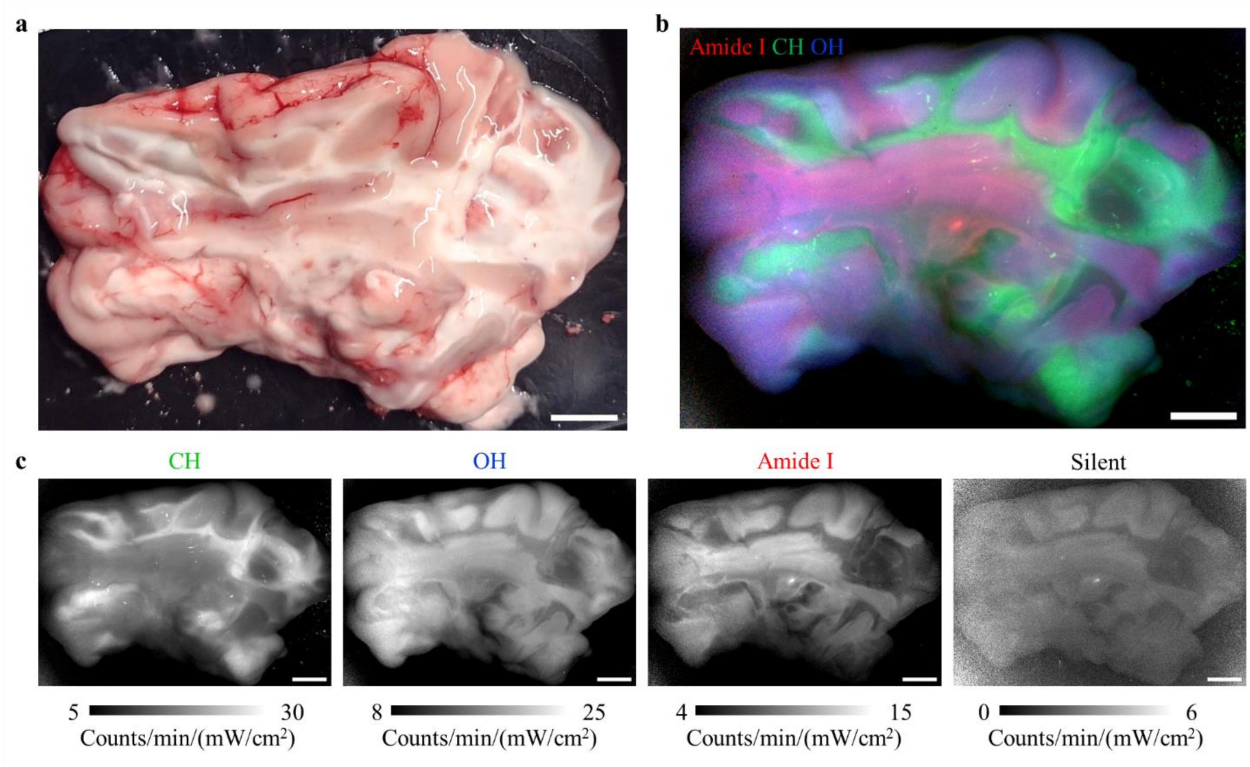

**Fig. S20.** Longitudinal section of the porcine left hemisphere (lateral view) showed in Fig. S19, imaged **(a)** in the visible and **(b-c)** using SWIR Raman imaging. Scale bar: 1 cm.

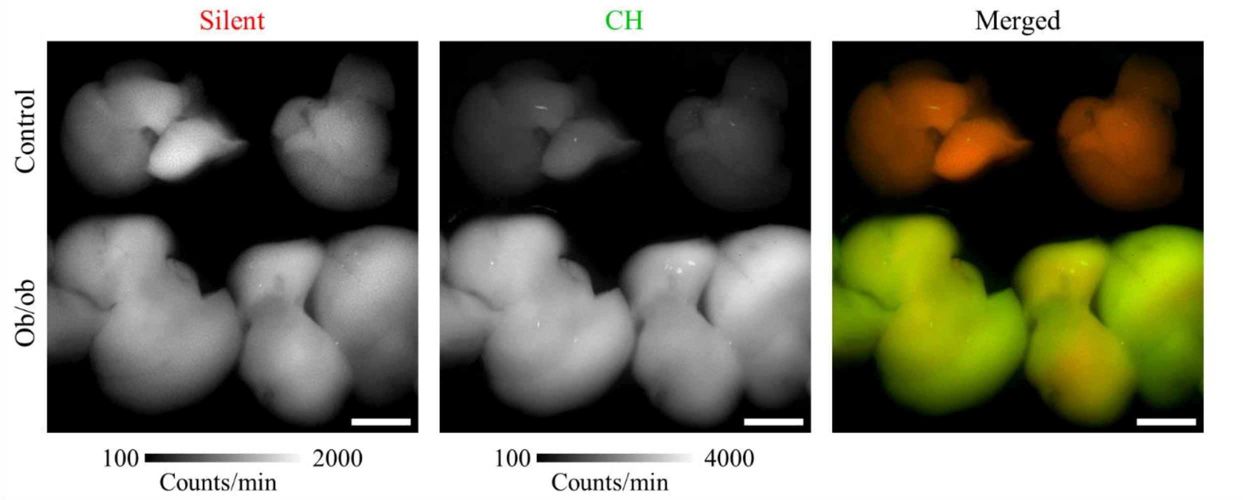

**Fig. S21**. Wide-field Raman imaging of livers excised from genetically obese mice (Ob/Ob) and lean C57BL/6J mice (Control). Two whole livers per group are shown. Scale bar: 1 cm.

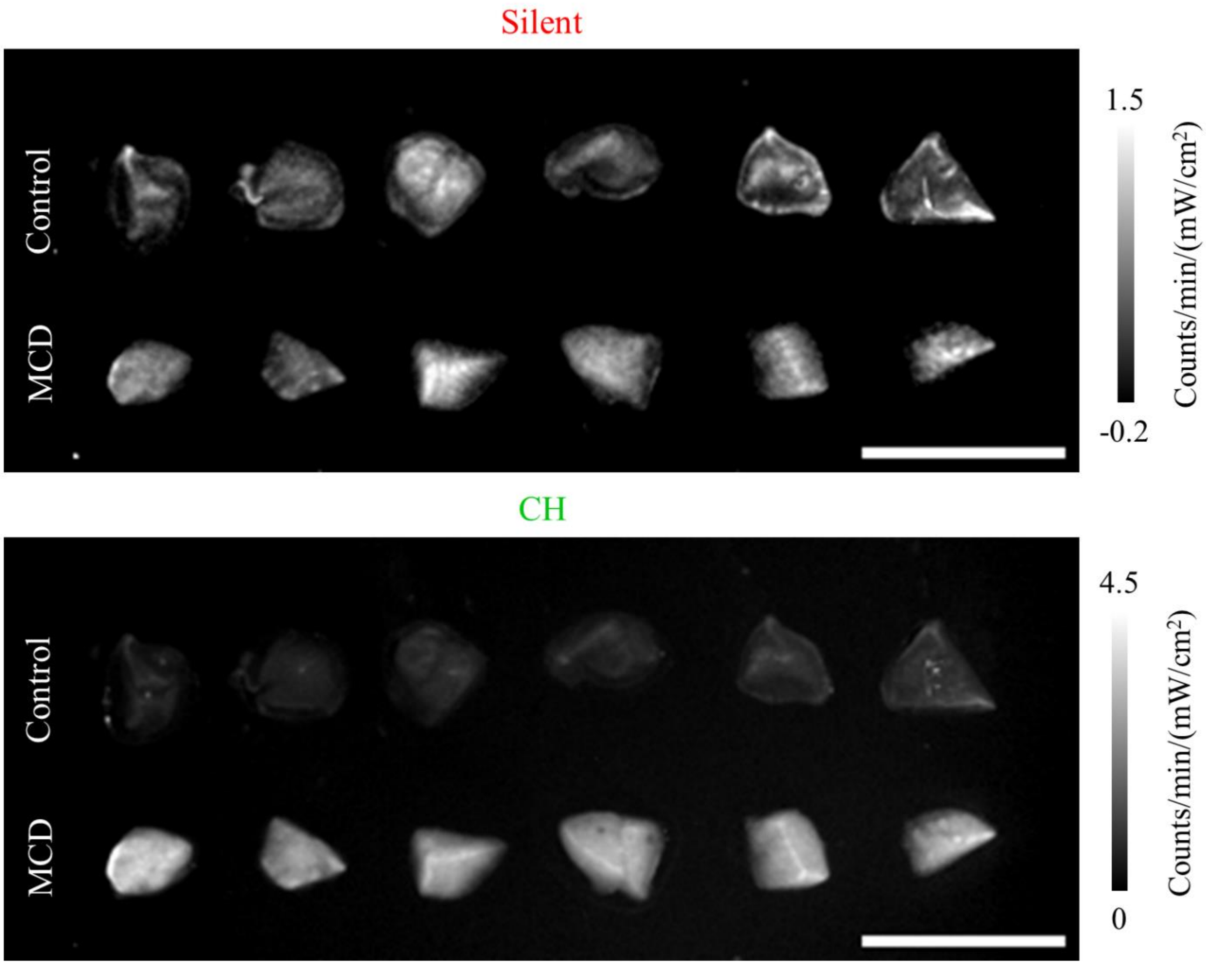

**Fig. S22**. Single-band images of the unfixed dissected livers pieces shown in Fig. 4 a. Samples from six mice per group are shown. MCD, methionine-choline diet. Scale bar: 1 cm.

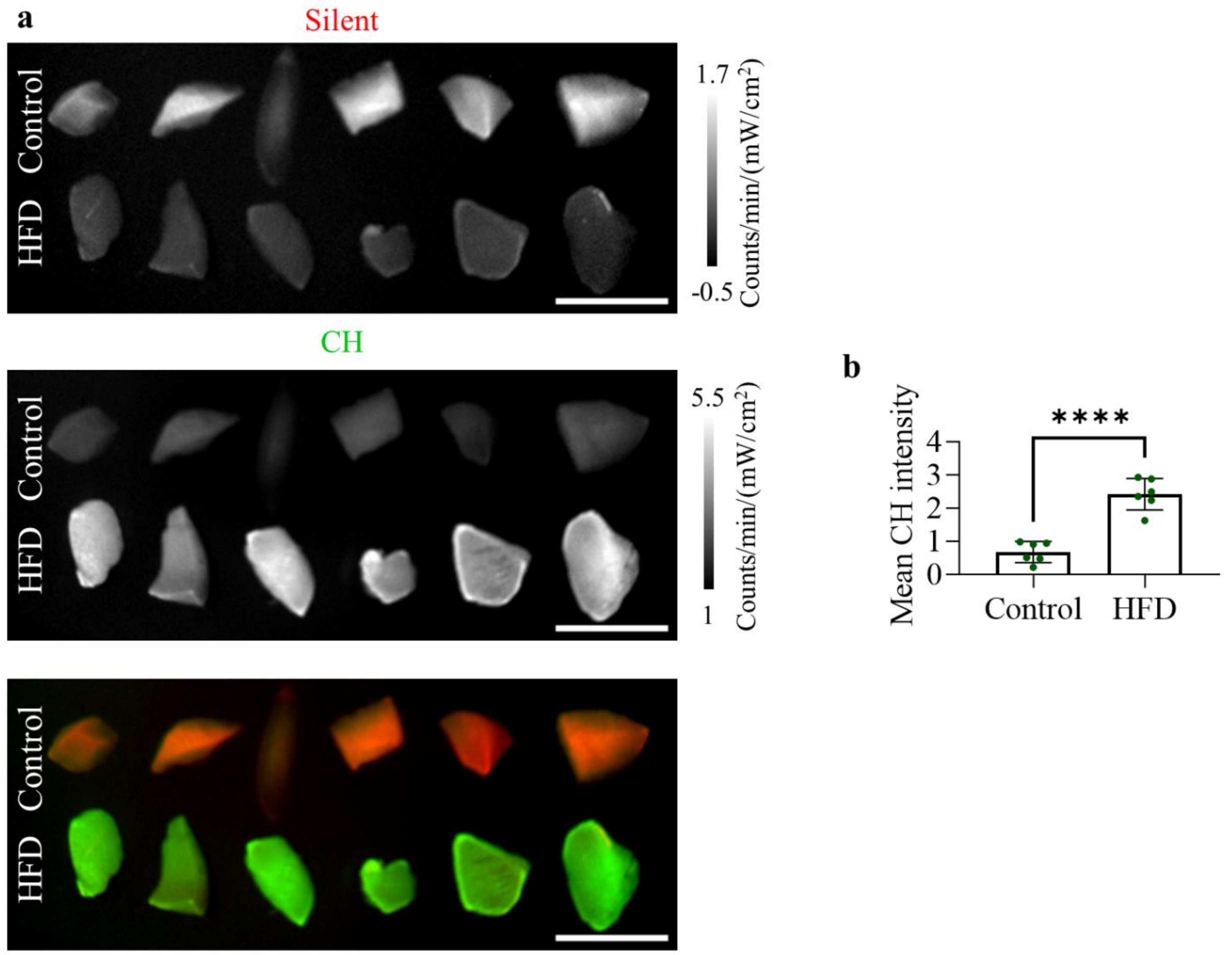

**Fig. S23**. Liver samples from mice induced either to a high fat diet (HFD) for 12 weeks or to a control diet. **a)** Samples illuminated at 1064 nm and imaged in the silent (red) and CH (green) Raman regions. **b)** Quantification of mean CH Raman intensity based on the imaging [counts/min/(mW/cm2)]. Differences between the groups were statistically analyzed using an unpaired, two-tailed t test. **** indicates P-value < 0.0001. Scale bar: 1 cm.

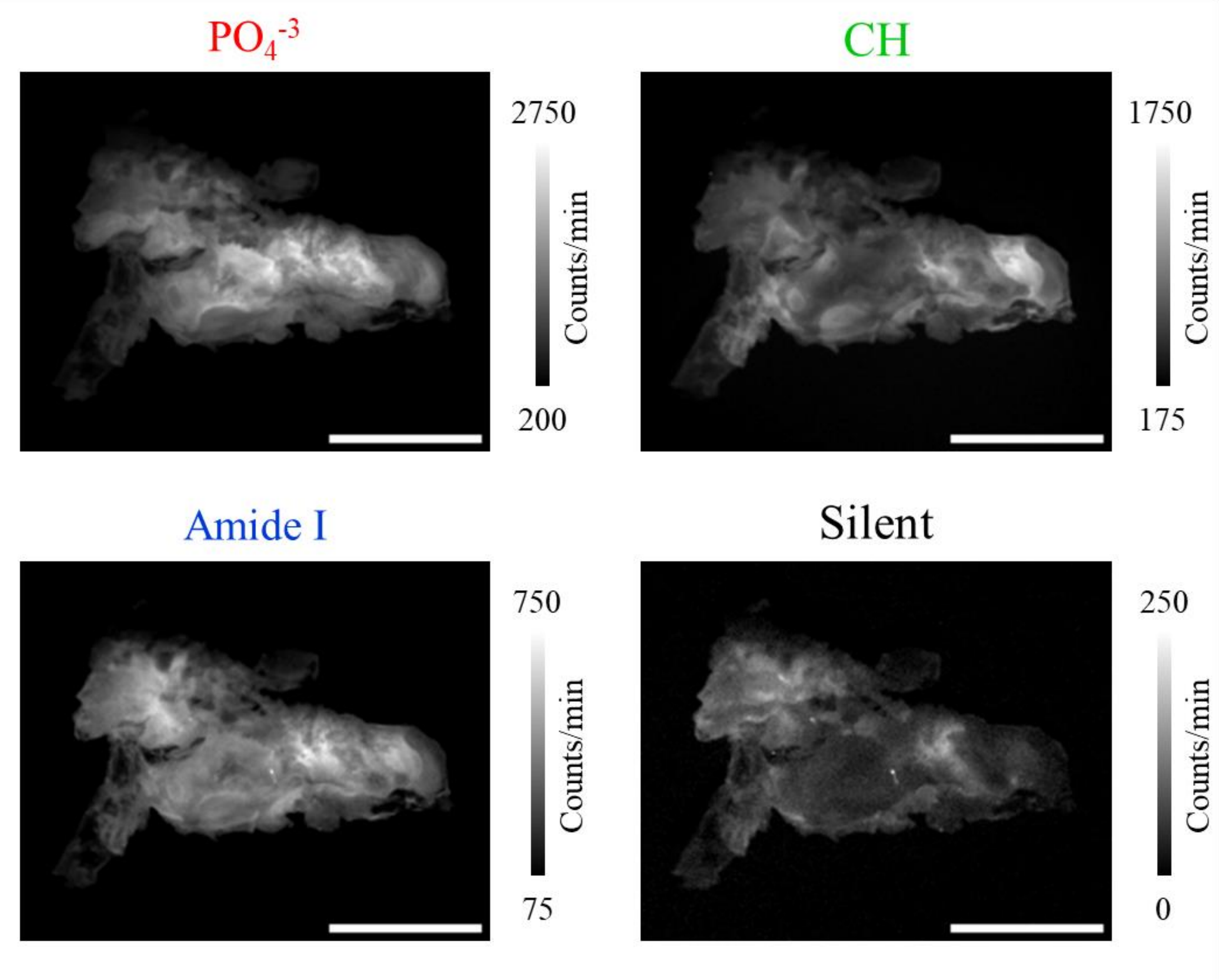

**Fig. S24**. Single-band images of the atherosclerotic plaque shown in Fig. 4 e, with the addition of an image of the silent region. Scale bar: 1 cm.

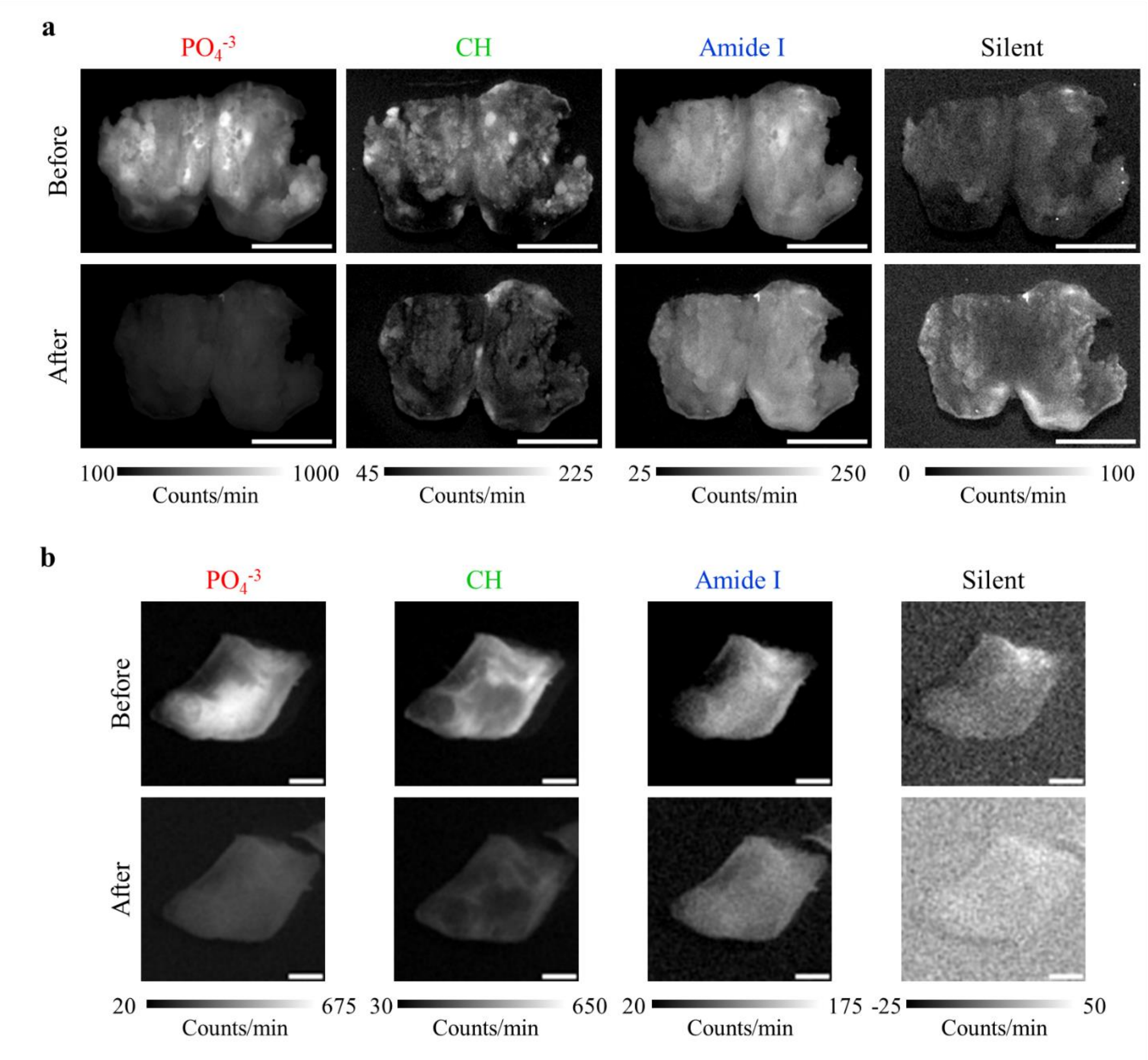

**Fig. S25**. Single-band images of the atherosclerotic plaques in (**a**) Fig. 4 f (femoral artery) and (**b**) in Fig. 4 g **(**carotid artery), with the addition of an image of the silent region. Scale bars: **(a)** 1 cm, **(b)** 0.2 cm.

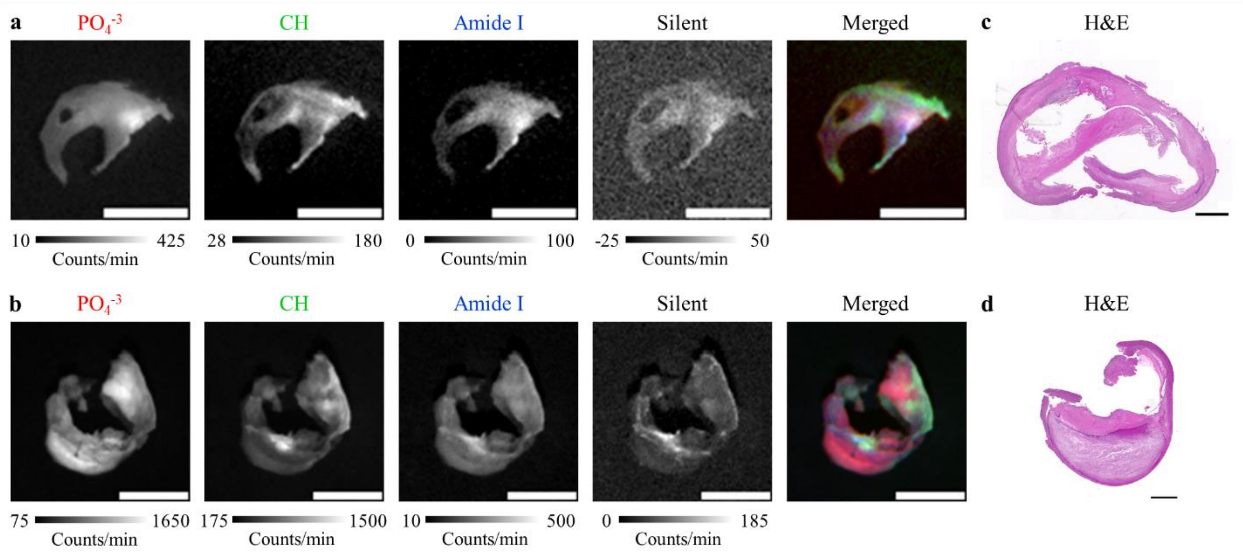

**Fig. S26**. **a-b)** Biopsied human atherosclerotic plaques from carotid arteries illuminated at 1064 nm and imaged in the $PO_4^{-3}$ (red), CH (green), and Amide I (blue) Raman bands. The silent region is also shown individually, but not in the merged image. **c-d**) Histological section stained with hematoxylin and eosin (H&E) adjacent to the unfixed samples shown in panels **(a)-(b)**. Scale bars: **(a)-(b)** 0.5 cm, **(c)-(d)** 0.1 cm.

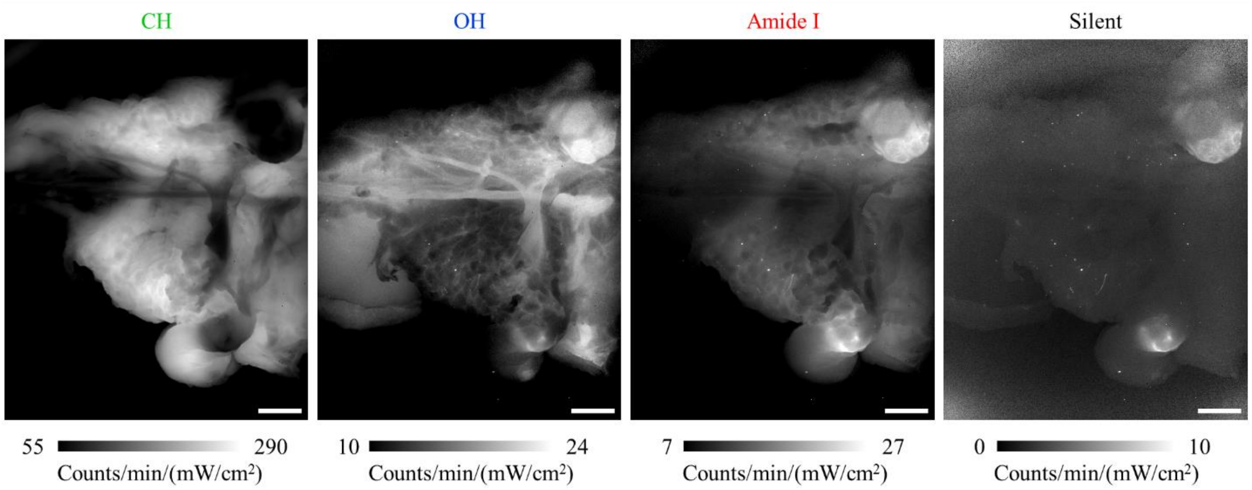

**Fig. S27**. Single-band images of the exposed porcine facial nerve shown in Fig. 5 b-c. Scale bar: 1 cm.

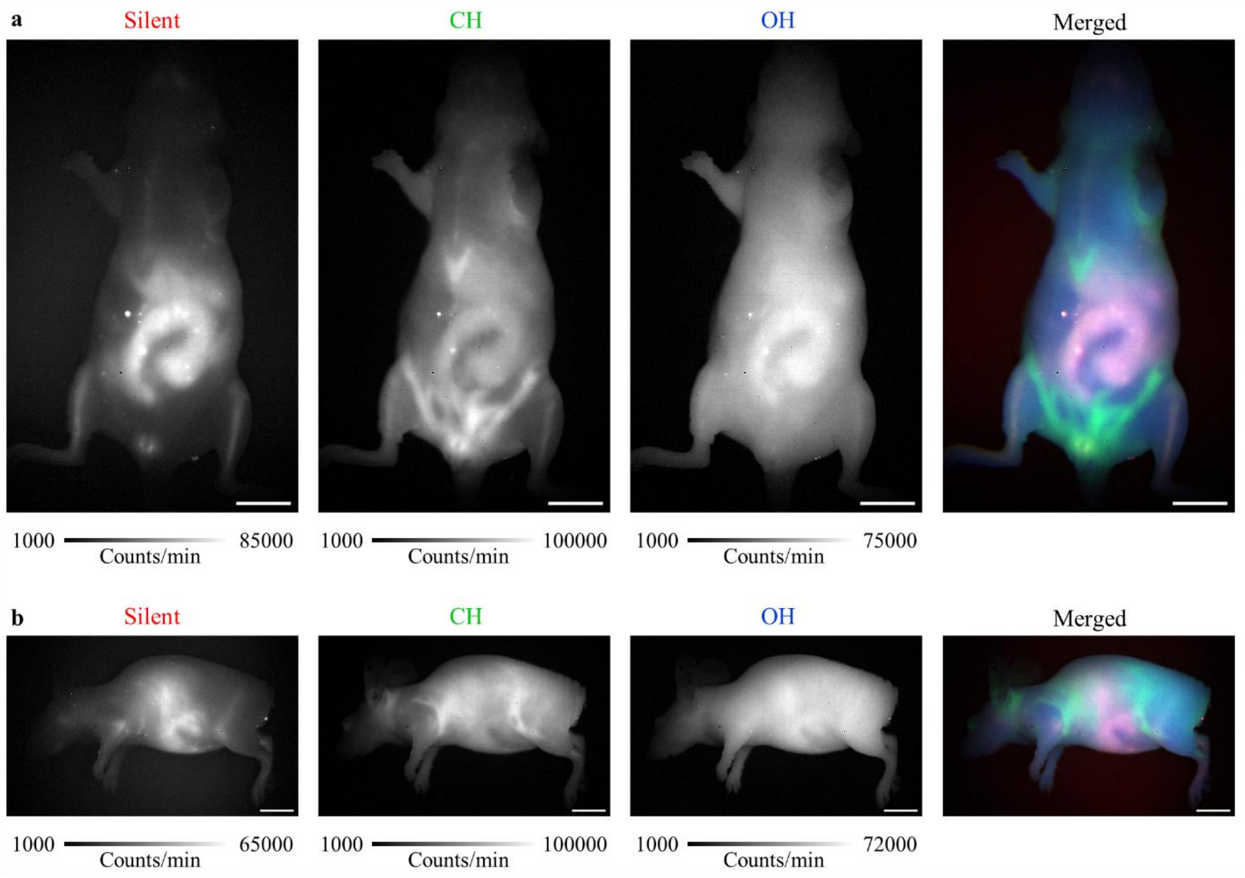

**Fig. S28**. (**a)** Ventral and (**b**) left lateral views of a mouse imaged with SWIR Raman imaging using a portable, air-cooled InGaAs detector upon 892 nm illumination. Scale bar: 1cm.

| Main Figure | Suppl. Figure | Band | Laser (nm) | Laser clean-up filter | Max laser pwr[1] | Exp. time (s) | Slot 1 (LP) | Filter slot 2 | Filter slot 3 | Slot 4 (LP) |
|---|---|---|---|---|---|---|---|---|---|---|
| Fig. 1 c | - | **CH** | 892 | 2xSP900 | 152 | 30 | 1100 | BP1200-25 | empty | 1100 |
| | | **OH** | 892 | 2xSP900 | 152 | 30 | 1100 | BP1300-25 | empty | 1100 |
| Fig. 1 d | S4 | **Silent** | 892 | 2xSP900 | 152 | 10 | 1100 | BP1150-25 | empty | 1100 |
| | | **CH** | 892 | 2xSP900 | 152 | 10 | 1100 | BP1200-25 | empty | 1100 |
| | | **OH** | 892 | 2xSP900 | 152 | 10 | 1100 | BP1300-25 | empty | 1100 |
| Fig. 1 e | S6 a | **CH** | 938 | SP1000 | 115 | 30 | BP1275-50 | LP1000* | BP1300-50 | 1100 |
| | | **OH** | 938 | SP1000 | 115 | 30 | BP1350-50 | LP1000* | empty | 1100 |
| | | **Silent** | 938 | SP1000 | 115 | 30 | BP1200-25 | LP1000* | empty | 1100 |
| Fig. 1 f | S6 b | **CH** | 938 | SP1000 | 115 | 10 | BP1275-50 | LP1000* | BP1300-50 | 1100 |
| | | **OH** | 938 | SP1000 | 115 | 10 | BP1350-50 | LP1000* | empty | 1100 |
| | | **Silent** | 938 | SP1000 | 115 | 10 | BP1200-25 | LP1000* | empty | 1100 |
| Fig. 2 a-b [785 nm] | S7 [785 nm] | **Silent** | 785 | SP0800 | 138 | 10 | 950 | BP950-50 | empty | 900 |
| | | **CH** | 785 | SP0800 | 138 | 10 | 950 | BP1000-50 | LP1000 | 1000 |
| | | **OH** | 785 | SP0800 | 138 | 10 | 950 | BP1050-50 | empty | 1050 |
| Fig. 2 a-b [892 nm] | S7 [892 nm] | **Silent** | 892 | SP900 | 145 | 30 | 950 | BP1150-25 | empty | 1100 |
| | | **CH** | 892 | SP900 | 145 | 30 | 950 | BP1200-25 | empty | 1100 |
| | | **OH** | 892 | SP900 | 145 | 30 | 950 | BP1300-25 | empty | 1100 |
| Fig. 2 a-b [938 nm] | S7 [938 nm] | **Silent** | 938 | SP950 | 153 | 30 | 1150 | BP1175-50 | empty | 1150 |
| | | **CH** | 938 | SP950 | 153 | 30 | 1150 | BP1300-50 | empty | 1150 |
| | | **OH** | 938 | SP950 | 153 | 30 | 1150 | BP1350-50 | empty | 1150 |
| Fig. 2 a-b [968 nm] | S7 [968 nm] | **Silent** | 968 | SP1000 | 152 | 45 | 1150 | BP1200-50 | empty | 1200 |
| | | **CH** | 968 | SP1000 | 152 | 45 | 1150 | BP1350-50 | empty | 1200 |
| | | **OH** | 968 | SP1000 | 152 | 45 | 1150 | BP1450-50 | empty | 1200 |
| Fig. 2 b [1064 nm] | | **Silent** | 1064 | SP1100 | 178 | 60 | 1150 | BP1400-50 | LP1100 | 1200 |

| | | | | | | | | | | |
|---|---|---|---|---|---|---|---|---|---|---|
| | **S7 [1064 nm]** | **CH** | 1064 | SP1100 | 178 | 60 | 1150 | BP1550-50 | LP1100 | 1200 |
| | | **OH** | 1064 | SP1100 | 178 | 60 | 1150 | BP1600-50 | LP1100 | 1200 |
| **Fig. 3 a** | **S10** | **Silent** | 892 | 2xSP900 | 152 | 30 | 1100 | BP1150-25 | empty | 1100 |
| | | **CH** | 892 | 2xSP900 | 152 | 30 | 1100 | BP1200-25 | empty | 1100 |
| | | **OH** | 892 | 2xSP900 | 152 | 30 | 1100 | BP1300-25 | empty | 1100 |
| **Fig. 3 b-c** | **S11** | **Silent** | 938 | SP950 | 159 | 45 | 1000* | BP1200-25 | LP1100 | 1150 |
| | | **CH** | 938 | SP950 | 159 | 45 | 1000* | BP1275-50 | BP1300-50 | 1150 |
| | | **OH** | 938 | SP950 | 159 | 45 | 1000* | BP1350-25 | LP1100 | 1150 |
| **Fig. 3 e [Before fasting]** | **S14** | **Silent** | 938 | SP950# | 143 | 30 | 1100 | BP1150-25 | BP1125-50 | 1100 |
| | | **CH** | 938 | SP950# | 143 | 30 | 1100 | BP1275-50 | BP1300-50 | 1100 |
| | | **OH** | 938 | SP950# | 143 | 30 | 1100 | BP1350-50 | BP1375-50 | 1100 |
| **Fig. 3 e [After fasting]** | **S14** | **Silent** | 938 | SP950# | 143 | 30 | 1100 | BP1150-25 | BP1125-50 | 1100 |
| | | **CH** | 938 | SP950# | 143 | 30 | 1100 | BP1275-50 | BP1300-50 | 1100 |
| | | **OH** | 938 | SP950# | 143 | 30 | 1100 | BP1350-50 | BP1375-50 | 1100 |
| **Fig. 3 e [24h after refeeding]** | **S14** | **Silent** | 938 | SP950# | 123 | 30 | 1100 | BP1150-25 | BP1125-50 | 1100 |
| | | **CH** | 938 | SP950# | 123 | 30 | 1100 | BP1275-50 | BP1300-50 | 1100 |
| | | **OH** | 938 | SP950# | 123 | 30 | 1100 | BP1350-50 | BP1375-50 | 1100 |
| **Fig. 3 e [48h after refeeding]** | **S14** | **Silent** | 938 | SP950# | 123 | 60 | 1100 | BP1150-25 | BP1125-50 | 1100 |
| | | **CH** | 938 | SP950# | 123 | 60 | 1100 | BP1275-50 | BP1300-50 | 1100 |
| | | **OH** | 938 | SP950# | 123 | 60 | 1100 | BP1350-50 | BP1375-50 | 1100 |
| **Fig. 4 a** | **S22** | **Silent** | 1064 | SP1100# | 115 | 600 | 2x1150 | BP1325-50 | BP1350-50 | 1250 |
| | | **CH** | 1064 | SP1100# | 115 | 600 | 2x1150 | BP1550-50 | empty | 1250 |
| **Fig. 4 e** | **S24** | **$PO4^{-3}$** | 1064 | SP1075# +BP1064-10 | 177 | 300 | 2x1150 | BP1175-50 | BP1200-50 | 1150 |
| | | **CH** | 1064 | SP1075# +BP1064-10 | 177 | 300 | 2x1150 | BP1550-50 | LP1200 | 1150 |
| | | **Amide I** | 1064 | SP1075# +BP1064-10 | 177 | 300 | 2x1150 | BP1275-50 | BP1300-50 | 1150 |

| | | | | | | | | | | |
|---|---|---|---|---|---|---|---|---|---|---|
| | | **Silent** | 1064 | SP1075# +BP1064-10 | 177 | 300 | 2x1150 | BP1350-50 | BP1375-50 | 1150 |
| **Fig. 4 f** | **S25 a** | **$PO4^{-3}$** | 1064 | SP1100+ SP1075# | 163 | 300 | WG81050 | BP1175-50 | BP1200-50 | 1150 |
| | | **CH** | 1064 | SP1100+ SP1075# | 163 | 300 | WG81050 | BP1550-50 | LP1150 | 1150 |
| | | **Amide I** | 1064 | SP1100+ SP1075# | 163 | 300 | WG81050 | BP1275-50 | BP1300-50 | 1150 |
| | | **Silent** | 1064 | SP1100+ SP1075# | 163 | 300 | WG81050 | BP1350-50 | BP1375-50 | 1150 |
| **Fig. 4 g** | **S25 b** | **$PO4^{-3}$** | 1064 | SP1100+ SP1075# | 163 | 300 | WG81050 | BP1175-50 | BP1200-50 | 1150 |
| | | **CH** | 1064 | SP1100+ SP1075# | 163 | 300 | WG81050 | BP1550-50 | LP1150 | 1150 |
| | | **Amide I** | 1064 | SP1100+ SP1075# | 163 | 300 | WG81050 | BP1275-50 | BP1300-50 | 1150 |
| | | **Silent** | 1064 | SP1100+ SP1075# | 163 | 300 | WG81050 | BP1350-50 | BP1375-50 | 1150 |
| **Fig. 5 b-c** | **S27** | **CH** | 938 | SP1000 | 115 | 60 | BP1275-50 | LP1000* | BP1300-50 | 1100 |
| | | **OH** | 938 | SP1000 | 115 | 60 | BP1350-50 | LP1000* | empty | 1100 |
| | | **Amide I** | 938 | SP1000 | 115 | 60 | BP1100-50 | LP1000* | BP1125-50 | 1100 |
| | | **Silent** | 938 | SP1000 | 115 | 60 | BP1200-25 | LP1000* | empty | 1100 |
| **-** | **S3** | **$PO4^{-3}$** | 1064 | SP1075# +BP1064-10 | 177 | 300 | 2x1150 | BP1275-50 | BP1300-50 | 1150 |
| | | **CH** | 1064 | SP1075# +BP1064-10 | 177 | 300 | 2x1150 | BP1550-50 | LP1200 | 1150 |
| | | **Amide I** | 1064 | SP1075# +BP1064-10 | 177 | 300 | 2x1150 | BP1175-50 | BP1200-50 | 1150 |
| | **S5** | **Silent** | 938 | SP950 | 153 | 30 | 1150 | BP1175-50 | empty | 1150 |
| | | **CH** | 938 | SP950 | 153 | 30 | 1150 | BP1300-50 | empty | 1150 |
| | | **OH** | 938 | SP950 | 153 | 30 | 1150 | BP1350-50 | empty | 1150 |
| | **S9 [785 nm]** | **-** | 785 | SP0800 | 138 | 10 | 950 | LP950 | empty | 900 |
| | **S9 [892 nm]** | **-** | 892 | SP900 | 145 | 30 | 950 | LP1100 | empty | 1100 |
| | **S9 [938 nm]** | **-** | 938 | SP950 | 153 | 30 | 1150 | LP1100 | empty | 1150 |

| | | | | | | | | | | |
|---|---|---|---|---|---|---|---|---|---|---|
| | **S9 [968 nm]** | **-** | 968 | SP1000 | 152 | 45 | 1150 | LP1100 | empty | 1200 |
| | **S9 [1064 nm]** | **-** | 1064 | SP1100 | 178 | 60 | 1150 | LP1350 | LP1100 | 1200 |
| | **S12** | **Silent** | 938 | SP950 | 159 | 45 | 1000* | BP1200-25 | LP1100 | 1150 |
| | | **CH** | 938 | SP950 | 159 | 45 | 1000* | BP1275-50 | BP1300-50 | 1150 |
| | | **OH** | 938 | SP950 | 159 | 45 | 1000* | BP1350-25 | LP1100 | 1150 |
| | **S15** | **Silent** | 892 | SP900 | 124 | 10 | 1100 | BP1150-25 | empty | 1100 |
| | | **CH** | 892 | SP900 | 124 | 10 | 1100 | BP1200-25 | empty | 1100 |
| | | **OH** | 892 | SP900 | 124 | 10 | 1100 | BP1300-25 | empty | 1100 |
| | **S16** | **Silent** | 892 | SP900 | 124 | 10 | 1100 | BP1150-25 | empty | 1100 |
| | | **CH** | 892 | SP900 | 124 | 10 | 1100 | BP1200-25 | empty | 1100 |
| | | **OH** | 892 | SP900 | 124 | 10 | 1100 | BP1300-25 | empty | 1100 |
| | **S17 [Days 3 and 5]** | **Silent** | 892 | SP900 | 106 | 20 | 1100 | BP1150-25 | empty | 1100 |
| | | **CH** | 892 | SP900 | 106 | 20 | 1100 | BP1200-25 | empty | 1100 |
| | | **OH** | 892 | SP900 | 106 | 20 | 1100 | BP1300-25 | empty | 1100 |
| | | **4T1-iRFP** | 672 | SP900 | 89 | 1 | 1100 | LP1100 | empty | 1100 |
| | **S17 [Days 10 and 16]** | **Silent** | 892 | SP900 | 104 | 20 | 1100 | BP1150-25 | empty | 1100 |
| | | **CH** | 892 | SP900 | 104 | 20 | 1100 | BP1200-25 | empty | 1100 |
| | | **OH** | 892 | SP900 | 104 | 20 | 1100 | BP1300-25 | empty | 1100 |
| | | **4T1-iRFP** | 672 | SP900 | 92 | 1 | 1100 | LP1100 | empty | 1100 |
| | **S18** | **Silent** | 892 | 2xSP900 | 172 | 30 | 950 | BP1150-25 | empty | 1100 |
| | | **CH** | 892 | 2xSP900 | 172 | 30 | 950 | BP1200-25 | empty | 1100 |
| | | **OH** | 892 | 2xSP900 | 172 | 30 | 950 | BP1300-25 | empty | 1100 |
| | **S19** | **CH** | 938 | SP1000 | 115 | 30 | BP1275-50 | LP1000* | BP1300-50 | 1100 |
| | | **OH** | 938 | SP1000 | 115 | 30 | BP1350-50 | LP1000* | empty | 1100 |
| | | **Amide I** | 938 | SP1000 | 115 | 30 | BP1100-50 | LP1000* | BP1125-50 | 1100 |
| | | **Silent** | 938 | SP1000 | 115 | 30 | BP1200-25 | LP1000* | empty | 1100 |

| | | | | | | | | | | |
|---|---|---|---|---|---|---|---|---|---|---|
| | **S20** | **CH** | 938 | SP1000 | 115 | 30 | BP1275-50 | LP1000* | BP1300-50 | 1100 |
| | | **OH** | 938 | SP1000 | 115 | 30 | BP1350-50 | LP1000* | empty | 1100 |
| | | **Amide I** | 938 | SP1000 | 115 | 30 | BP1100-50 | LP1000* | BP1125-50 | 1100 |
| | | **Silent** | 938 | SP1000 | 115 | 30 | BP1200-25 | LP1000* | empty | 1100 |
| | **S21** | **Silent** | 938 | SP950 | 96 | 60 | 1000* | BP1125-50 | BP1150-25 | 1100 |
| | | **CH** | 938 | SP950 | 96 | 60 | 1000* | BP1275-50 | BP1300-50 | 1100 |
| | **S23** | **Silent** | 1064 | SP1100# | 115 | 600 | 2x1150 | BP1325-50 | BP1350-50 | 1250 |
| | | **CH** | 1064 | SP1100# | 115 | 600 | 2x1150 | BP1550-50 | empty | 1250 |
| | **S26 a** | **PO4$^{-3}$** | 1064 | SP1100+ SP1075# | 163 | 300 | WG81050 | BP1175-50 | BP1200-50 | 1150 |
| | | **CH** | 1064 | SP1100+ SP1075# | 163 | 300 | WG81050 | BP1550-50 | LP1150 | 1150 |
| | | **Amide I** | 1064 | SP1100+ SP1075# | 163 | 300 | WG81050 | BP1275-50 | BP1300-50 | 1150 |
| | | **Silent** | 1064 | SP1100+ SP1075# | 163 | 300 | WG81050 | BP1350-50 | BP1375-50 | 1150 |
| | **S26 c** | **PO4$^{-3}$** | 1064 | SP1100+ SP1075# | 176 | 300 | 2x1150 | BP1175-50 | BP1200-50 | 1150 |
| | | **CH** | 1064 | SP1100+ SP1075# | 176 | 300 | 2x1150 | BP1550-50 | LP1150 | 1150 |
| | | **Amide I** | 1064 | SP1100+ SP1075# | 176 | 300 | 2x1150 | BP1275-50 | BP1300-50 | 1150 |
| | | **Silent** | 1064 | SP1100+ SP1075# | 176 | 300 | 2x1150 | BP1350-50 | BP1375-50 | 1150 |
| | **S28** | **Silent** | 892 | SP900 | 159 | 7.5 | 1100 | BP1150-50 | empty | 1100 |
| | | **CH** | 892 | SP900 | 159 | 7.5 | 1100 | BP1200-50 | empty | 1100 |
| | | **OH** | 892 | SP900 | 159 | 7.5 | 1100 | BP1300-50 | empty | 1100 |

**Table S1**. SWIR Raman acquisition configurations for each imaging band shown in the figures. Band-pass filters (BP) are denominated as central wavelength (nm) – width (nm). LP, long-pass filter; SP, short-pass filter; WG, WG81050 (Thorlabs); Exp. time, exposure time, Max laser pwr[1], maximum laser power density in the field of view, in mW/cm². All SP laser clean-up filters are from the Thorlabs FESH series, except for the ones labeled with "#", which were purchased from Edmund Optics (all 1-in filters). BP1064-10 is catalog no. FLH1064-10 (Thorlabs). All LP filters from slots 1 to 3 are from the Thorlabs FELH series, except for the ones labeled with “*”, which were purchased from Edmund Optics (all 1-in filters). All Slot 4 filters are 2-in filters from Edmund Optics. All BP filters are from Edmund Optics.

| Main Figure | Suppl. Figure | Band | Laser (nm) | Effective Emission Filter | Effective Filtered Range (wavelength, nm) | Effective Filtered Range (wavenumber, $cm^{-1}$) |
|---|---|---|---|---|---|---|
| **Fig. 1 c** | **-** | **CH** | 892 | BP1200.0-25.0 | 1187.5 - 1212.5 | 2789.7 - 2963.3 |
| | | **OH** | 892 | BP1300.0-25.0 | 1287.5 - 1312.5 | 3443.8 - 3591.7 |
| **Fig. 1 d** | **S4** | **Silent** | 892 | BP1150.0-25.0 | 1137.5 - 1162.5 | 2419.6 - 2608.6 |
| | | **CH** | 892 | BP1200.0-25.0 | 1187.5 - 1212.5 | 2789.7 - 2963.3 |
| | | **OH** | 892 | BP1300.0-25.0 | 1287.5 - 1312.5 | 3443.8 - 3591.7 |
| **Fig. 1 e** | **S6 a** | **CH** | 938 | BP1287.5-25.0 | 1275.0 - 1300.0 | 2817.8 - 2968.7 |
| | | **OH** | 938 | BP1350.0-50.0 | 1325.0 - 1375.0 | 3113.8 - 3388.3 |
| | | **Silent** | 938 | BP1200.0-25.0 | 1187.5 - 1212.5 | 2239.9 - 2413.6 |
| **Fig. 1 f** | **S6 b** | **CH** | 938 | BP1287.5-25.0 | 1275.0 - 1300.0 | 2817.8 - 2968.7 |
| | | **OH** | 938 | BP1350.0-50.0 | 1325.0 - 1375.0 | 3113.8 - 3388.3 |
| | | **Silent** | 938 | BP1200.0-25.0 | 1187.5 - 1212.5 | 2239.9 - 2413.6 |
| **Fig. 2 a-b [785 nm]** | **S7 [785 nm]** | **Silent** | 785 | BP962.5-25.0 | 950.0 - 975.0 | 2212.5 - 2482.4 |
| | | **CH** | 785 | BP1012.5-25.0 | 1000.0 - 1025.0 | 2738.9 - 2982.8 |
| | | **OH** | 785 | BP1062.5-25.0 | 1050.0 - 1075.0 | 3215.0 - 3436.5 |
| **Fig. 2 a-b [892 nm]** | **S7 [892 nm]** | **Silent** | 892 | BP1150.0-25.0 | 1137.5 - 1162.5 | 2419.6 - 2608.6 |
| | | **CH** | 892 | BP1200.0-25.0 | 1187.5 - 1212.5 | 2789.7 - 2963.3 |
| | | **OH** | 892 | BP1300.0-25.0 | 1287.5 - 1312.5 | 3443.8 - 3591.7 |
| **Fig. 2 a-b [938 nm]** | **S7 [938 nm]** | **Silent** | 938 | BP1175.0-50.0 | 1150.0 - 1200.0 | 1965.3 - 2327.6 |
| | | **CH** | 938 | BP1300.0-50.0 | 1275.0 - 1325.0 | 2817.8 - 3113.8 |
| | | **OH** | 938 | BP1350.0-50.0 | 1325.0 - 1375.0 | 3113.8 - 3388.3 |
| **Fig. 2 a-b [968 nm]** | **S7 [968 nm]** | **Silent** | 968 | BP1212.5-25.0 | 1200.0 - 1225.0 | 1997.2 - 2167.3 |
| | | **CH** | 968 | BP1350.0-50.0 | 1325.0 - 1375.0 | 2783.4 - 3057.9 |
| | | **OH** | 968 | BP1450.0-50.0 | 1425.0 - 1475.0 | 3313.0 - 3550.9 |
| **Fig. 2 b [1064 nm]** | **S7 [1064 nm]** | **Silent** | 1064 | BP1400.0-50.0 | 1375.0 - 1425.0 | 2125.8 - 2381.0 |
| | | **CH** | 1064 | BP1550.0-50.0 | 1525.0 - 1575.0 | 2841.1 - 3049.3 |
| | | **OH** | 1064 | BP1600.0-50.0 | 1575.0 - 1625.0 | 3049.3 - 3244.7 |
| **Fig. 3 a** | **S10** | **Silent** | 892 | BP1150.0-25.0 | 1137.5 - 1162.5 | 2419.6 - 2608.6 |
| | | **CH** | 892 | BP1200.0-25.0 | 1187.5 - 1212.5 | 2789.7 - 2963.3 |
| | | **OH** | 892 | BP1300.0-25.0 | 1287.5 - 1312.5 | 3443.8 - 3591.7 |
| **Fig. 3 b-c** | **S11** | **Silent** | 938 | BP1200.0-25.0 | 1187.5 - 1212.5 | 2239.9 - 2413.6 |
| | | **CH** | 938 | BP1287.5-25.0 | 1275.0 - 1300.0 | 2817.8 - 2968.7 |
| | | **OH** | 938 | BP1350.0-25.0 | 1337.5 - 1362.5 | 3184.3 - 3321.5 |
| **Fig. 3 e [Before fasting]** | **S14** | **Silent** | 938 | BP1143.75-12.5 | 1137.5 - 1150.0 | 1869.8 - 1965.3 |
| | | **CH** | 938 | BP1287.5-25.0 | 1275.0 - 1300.0 | 2817.8 - 2968.7 |
| | | **OH** | 938 | BP1362.5-25.0 | 1350.0 - 1375.0 | 3253.6 - 3388.3 |

| | | | | | | |
|---|---|---|---|---|---|---|
| **Fig. 3 e [After fasting]** | **S14** | **Silent** | 938 | BP1143.75-12.5 | 1137.5 - 1150.0 | 1869.8 - 1965.3 |
| | | **CH** | 938 | BP1287.5-25.0 | 1275.0 - 1300.0 | 2817.8 - 2968.7 |
| | | **OH** | 938 | BP1362.5-25.0 | 1350.0 - 1375.0 | 3253.6 - 3388.3 |
| **Fig. 3 e [24h after refeeding]** | **S14** | **Silent** | 938 | BP1143.75-12.5 | 1137.5 - 1150.0 | 1869.8 - 1965.3 |
| | | **CH** | 938 | BP1287.5-25.0 | 1275.0 - 1300.0 | 2817.8 - 2968.7 |
| | | **OH** | 938 | BP1362.5-25.0 | 1350.0 - 1375.0 | 3253.6 - 3388.3 |
| **Fig. 3 e [48h after refeeding]** | **S14** | **Silent** | 938 | BP1143.75-12.5 | 1137.5 - 1150.0 | 1869.8 - 1965.3 |
| | | **CH** | 938 | BP1287.5-25.0 | 1275.0 - 1300.0 | 2817.8 - 2968.7 |
| | | **OH** | 938 | BP1362.5-25.0 | 1350.0 - 1375.0 | 3253.6 - 3388.3 |
| **Fig. 4 a** | **S22** | **Silent** | 1064 | BP1337.5-25.0 | 1325.0 - 1350.0 | 1851.3 - 1991.1 |
| | | **CH** | 1064 | BP1550.0-50.0 | 1525.0 - 1575.0 | 2841.1 - 3049.3 |
| **Fig. 4 e** | **S24** | **$PO4^{-3}$** | 1064 | BP1187.5-25.0 | 1175.0 - 1200.0 | 887.9 - 1065.2 |
| | | **CH** | 1064 | BP1550.0-50.0 | 1525.0 - 1575.0 | 2841.1 - 3049.3 |
| | | **Amide I** | 1064 | BP1287.5-25.0 | 1275.0 - 1300.0 | 1555.4 - 1706.2 |
| | | **Silent** | 1064 | BP1362.5-25.0 | 1350.0 - 1375.0 | 1991.1 - 2125.8 |
| **Fig. 4 f** | **S25 a** | **$PO4^{-3}$** | 1064 | BP1187.5-25.0 | 1175.0 - 1200.0 | 887.9 - 1065.2 |
| | | **CH** | 1064 | BP1550.0-50.0 | 1525.0 - 1575.0 | 2841.1 - 3049.3 |
| | | **Amide I** | 1064 | BP1287.5-25.0 | 1275.0 - 1300.0 | 1555.4 - 1706.2 |
| | | **Silent** | 1064 | BP1362.5-25.0 | 1350.0 - 1375.0 | 1991.1 - 2125.8 |
| **Fig. 4 g** | **S25 b** | **$PO4^{-3}$** | 1064 | BP1187.5-25.0 | 1175.0 - 1200.0 | 887.9 - 1065.2 |
| | | **CH** | 1064 | BP1550.0-50.0 | 1525.0 - 1575.0 | 2841.1 - 3049.3 |
| | | **Amide I** | 1064 | BP1287.5-25.0 | 1275.0 - 1300.0 | 1555.4 - 1706.2 |
| | | **Silent** | 1064 | BP1362.5-25.0 | 1350.0 - 1375.0 | 1991.1 - 2125.8 |
| **Fig. 5 b-c** | **S27** | **CH** | 938 | BP1287.5-25.0 | 1275.0 - 1300.0 | 2817.8 - 2968.7 |
| | | **OH** | 938 | BP1350.0-50.0 | 1325.0 - 1375.0 | 3113.8 - 3388.3 |
| | | **Amide I** | 938 | BP1112.5-25.0 | 1100.0 - 1125.0 | 1570.1 - 1772.1 |
| | | **Silent** | 938 | BP1200.0-25.0 | 1187.5 - 1212.5 | 2239.9 - 2413.6 |
| **-** | **S3** | **$PO4^{-3}$** | 1064 | BP1287.5-25.0 | 1275.0 - 1300.0 | 1555.4 - 1706.2 |
| | | **CH** | 1064 | BP1550.0-50.0 | 1525.0 - 1575.0 | 2841.1 - 3049.3 |
| | | **Amide I** | 1064 | BP1187.5-25.0 | 1175.0 - 1200.0 | 887.9 - 1065.2 |
| | **S5** | **Silent** | 938 | BP1175.0-50.0 | 1150.0 - 1200.0 | 1965.3 - 2327.6 |
| | | **CH** | 938 | BP1300.0-50.0 | 1275.0 - 1325.0 | 2817.8 - 3113.8 |
| | | **OH** | 938 | BP1350.0-50.0 | 1325.0 - 1375.0 | 3113.8 - 3388.3 |
| | **S9 [785 nm]** | **-** | 785 | LP950 | 950.0 - 1600 | 2212.5 - 6488.9 |
| | **S9 [892 nm]** | **-** | 892 | LP1100 | 1100.0 - 1600 | 2119.9 - 4960.8 |
| | **S9 [938 nm]** | **-** | 938 | LP1150 | 1150.0 - 1600 | 1965.3 - 4411.0 |

| | | | | | | |
|---|---|---|---|---|---|---|
| | **S9 [968 nm]** | **-** | 968 | LP1200 | 1200.0 - 1600 | 1997.2 - 4080.6 |
| | **S9 [1064 nm]** | **-** | 1064 | LP1350 | 1350.0 - 1600 | 1991.1 - 3148.5 |
| | **S12** | **Silent** | 938 | BP1200.0-25.0 | 1187.5 - 1212.5 | 2239.9 - 2413.6 |
| | | **CH** | 938 | BP1287.5-25.0 | 1275.0 - 1300.0 | 2817.8 - 2968.7 |
| | | **OH** | 938 | BP1350.0-25.0 | 1337.5 - 1362.5 | 3184.3 - 3321.5 |
| | **S15** | **Silent** | 892 | BP1150.0-25.0 | 1137.5 - 1162.5 | 2419.6 - 2608.6 |
| | | **CH** | 892 | BP1200.0-25.0 | 1187.5 - 1212.5 | 2789.7 - 2963.3 |
| | | **OH** | 892 | BP1300.0-25.0 | 1287.5 - 1312.5 | 3443.8 - 3591.7 |
| | **S16** | **Silent** | 892 | BP1150.0-25.0 | 1137.5 - 1162.5 | 2419.6 - 2608.6 |
| | | **CH** | 892 | BP1200.0-25.0 | 1187.5 - 1212.5 | 2789.7 - 2963.3 |
| | | **OH** | 892 | BP1300.0-25.0 | 1287.5 - 1312.5 | 3443.8 - 3591.7 |
| | **S17 [Days 3 and 5]** | **Silent** | 892 | BP1150.0-25.0 | 1137.5 - 1162.5 | 2419.6 - 2608.6 |
| | | **CH** | 892 | BP1200.0-25.0 | 1187.5 - 1212.5 | 2789.7 - 2963.3 |
| | | **OH** | 892 | BP1300.0-25.0 | 1287.5 - 1312.5 | 3443.8 - 3591.7 |
| | | **4T1-iRFP** | 672 | LP1100 | 1100.0 - 1600 | 5790.0 - 8631.0 |
| | **S17 [Days 10 and 16]** | **Silent** | 892 | BP1150.0-25.0 | 1137.5 - 1162.5 | 2419.6 - 2608.6 |
| | | **CH** | 892 | BP1200.0-25.0 | 1187.5 - 1212.5 | 2789.7 - 2963.3 |
| | | **OH** | 892 | BP1300.0-25.0 | 1287.5 - 1312.5 | 3443.8 - 3591.7 |
| | | **4T1-iRFP** | 672 | LP1100 | 1100.0 - 1600 | 5790.0 - 8631.0 |
| | **S18** | **Silent** | 892 | BP1150.0-25.0 | 1137.5 - 1162.5 | 2419.6 - 2608.6 |
| | | **CH** | 892 | BP1200.0-25.0 | 1187.5 - 1212.5 | 2789.7 - 2963.3 |
| | | **OH** | 892 | BP1300.0-25.0 | 1287.5 - 1312.5 | 3443.8 - 3591.7 |
| | **S19** | **CH** | 938 | BP1287.5-25.0 | 1275.0 - 1300.0 | 2817.8 - 2968.7 |
| | | **OH** | 938 | BP1350.0-50.0 | 1325.0 - 1375.0 | 3113.8 - 3388.3 |
| | | **Amide I** | 938 | BP1112.5-25.0 | 1100.0 - 1125.0 | 1570.1 - 1772.1 |
| | | **Silent** | 938 | BP1200.0-25.0 | 1187.5 - 1212.5 | 2239.9 - 2413.6 |
| | **S20** | **CH** | 938 | BP1287.5-25.0 | 1275.0 - 1300.0 | 2817.8 - 2968.7 |
| | | **OH** | 938 | BP1350.0-50.0 | 1325.0 - 1375.0 | 3113.8 - 3388.3 |
| | | **Amide I** | 938 | BP1112.5-25.0 | 1100.0 - 1125.0 | 1570.1 - 1772.1 |
| | | **Silent** | 938 | BP1200.0-25.0 | 1187.5 - 1212.5 | 2239.9 - 2413.6 |
| | **S21** | **Silent** | 938 | BP1143.75-12.5 | 1137.5 - 1150.0 | 1869.8 - 1965.3 |
| | | **CH** | 938 | BP1287.5-25.0 | 1275.0 - 1300.0 | 2817.8 - 2968.7 |
| | **S23** | **Silent** | 1064 | BP1337.5-25.0 | 1325.0 - 1350.0 | 1851.3 - 1991.1 |
| | | **CH** | 1064 | BP1550.0-50.0 | 1525.0 - 1575.0 | 2841.1 - 3049.3 |
| | **S26 a** | **$PO4^{-3}$** | 1064 | BP1187.5-25.0 | 1175.0 - 1200.0 | 887.9 - 1065.2 |
| | | **CH** | 1064 | BP1550.0-50.0 | 1525.0 - 1575.0 | 2841.1 - 3049.3 |
| | | **Amide I** | 1064 | BP1287.5-25.0 | 1275.0 - 1300.0 | 1555.4 - 1706.2 |
| | | **Silent** | 1064 | BP1362.5-25.0 | 1350.0 - 1375.0 | 1991.1 - 2125.8 |

| | | | | | | |
|---|---|---|---|---|---|---|
| | **S26 c** | **$PO4^{-3}$** | 1064 | BP1187.5-25.0 | 1175.0 - 1200.0 | 887.9 - 1065.2 |
| | | **CH** | 1064 | BP1550.0-50.0 | 1525.0 - 1575.0 | 2841.1 - 3049.3 |
| | | **Amide I** | 1064 | BP1287.5-25.0 | 1275.0 - 1300.0 | 1555.4 - 1706.2 |
| | | **Silent** | 1064 | BP1362.5-25.0 | 1350.0 - 1375.0 | 1991.1 - 2125.8 |
| | **S28** | **Silent** | 892 | BP1150.0-50.0 | 1125.0 - 1175.0 | 2321.9 - 2700.1 |
| | | **CH** | 892 | BP1200.0-50.0 | 1175.0 - 1225.0 | 2700.1 - 3047.5 |
| | | **OH** | 892 | BP1300.0-50.0 | 1275.0 - 1325.0 | 3367.6 - 3663.6 |

**Table S2**. Laser and effective filter spectral range combination to achieve the specific SWIR Raman imaging bands for each figure. Effective emission filters were calculated from the combination of the filters in Slots 1 to 4 (Table S1). Ranges are represented as minimum – maximum of wavelengths or wavenumbers for each effective emission filter. The "effective filtered range (wavelength, nm)" uses the central wavelength (nm) – width (nm) of band-pass (BP) filters, when applicable, and 1600 nm as the detection limit of our camera. The "effective filtered range (wavenumber, $cm^{-1}$)" corresponds to the conversion of these detection ranges to wavenumbers for each particular illumination wavelength, according to Equation 1.
